# Evaluating acidotropic dyes for detecting mixotrophy in protists: Insights from cultures and field communities

**DOI:** 10.1101/2025.09.29.679303

**Authors:** Claire C. Z. Cook, Erica M. Ewton, Adrian Marchetti, Susanne Menden-Deuer, Nicole C. Millette, Shai Slomka, Emily V. Speciale, Susanne Wilken, Natalie R. Cohen

## Abstract

Mixotrophic protists combine photoautotrophic primary production with heterotrophic phagotrophy, and distinctly impact nutrient cycling and microbial food web dynamics in aquatic environments. Despite their biogeochemical importance, detecting and quantifying mixotrophic presence and grazing *in situ* remains challenging, preventing a comprehensive understanding of their ecology and biogeography. Fluorescently labeled particle (FLP) incubations are commonly used to quantify mixotroph abundance and ingestion but may underestimate activity due to prey and size preferences of grazers. Acidotropic dyes that stain acidic vacuoles associated with phagotrophy have emerged as an alternative to FLP incubations for estimating mixotroph abundance, yet have not been thoroughly tested among a diverse suite of marine eukaryotes. Here, we evaluate the effectiveness and specificity of two dyes, LysoTracker Green and LysoSensor Blue, in laboratory cultures and natural marine communities. In laboratory cultures, both dyes correctly did not stain one photoautotrophic species. However, LysoSensor failed to stain several known mixotrophs, indicating false negatives, while both dyes stained photoautotrophic diatoms, indicating false positives. In the field, LysoTracker staining broadly tracked with FLP-derived results in the North East Shelf (NES) and the diatom-rich California Current System (CCS). Both methods indicated lower mixotroph abundance and proportion in the CCS, suggesting acidotropic dyes may more reliably reflect mixotrophy in the field than in monoculture. This study highlights the utility and limitations of acidotropic dyes for detecting mixotrophy and underscores the importance of incorporating community composition and complementary grazing estimates for reliable interpretation.

## Introduction

The microbial loop plays a vital role in oceanic carbon and nutrient cycling. Advancements in the understanding of the microbial loop have emphasized the importance of mixotrophic organisms, which combine photosynthesis and heterotrophic consumption of prey within a single cell (Flynn et al., 2012; Glibert & Mitra, 2022; Mitra et al., 2016; Stoecker et al., 2017). Mixotrophs are hypothesized to play a crucial role in marine ecosystems by enhancing trophic transfer efficiency and recycling of inorganic nutrients compared to trophic specialists (Stoecker et al., 2017; Ward & Follows, 2016). Quantifying global distributions of mixotrophs will improve predictions of their role in ocean nutrient cycling and how changing ocean conditions will affect global biogeochemical processes such as carbon flow and nutrient dynamics (Millette et al., 2023).

Identifying mixotrophs *in situ* remains difficult despite the increasingly recognized ecological importance of these organisms within the global ocean (Beisner et al., 2019; Millette et al., 2023). Traditional methods to quantify plankton abundance, such as chlorophyll presence or grazing assays, were developed to target only photoautotrophs or heterotrophs, excluding mixotrophs that employ both trophic modes. Historically, the primary method for detecting mixotrophs has been fluorescently labeled particle (FLP) incubations, which rely on mixotrophs ingesting labeled particles over a short (minutes to hours) period of time (Bird & Kalff, 1986; Christaki et al., 1999; Li et al., 2023; McKie-Krisberg et al., 2015; Millette et al., 2017; Sanders & Porter, 1988; Unrein et al., 2007). As mixotrophic organisms may selectively not feed on surrogate prey due to particle size or artificial properties compared to natural prey, this approach is limited and likely underestimates mixotroph populations in natural systems (Stoecker et al., 2017; Unrein et al., 2007). The short incubation time of FLP incubations also captures behavior during one snapshot in time, potentially missing diel phagotrophic behavior occurring outside of incubation periods. Furthermore, microscopy or flow cytometry is typically used to quantify consumption of the fluorescent surrogate prey, and both analysis methods have drawbacks. Microscopy inherently introduces human error as the operator is required to distinguish between prey that is consumed versus that on the outside of the cell, and low fluorescence features may be missed. Flow cytometry demonstrates a similar issue with the potential for particles to be stuck on the outside of cells, resulting in false instances of ingestion. Thus, FLP incubations have caveats, and complementary methods for measuring mixotrophy would be valuable.

Acidotropic dyes, such as LysoTracker and LysoSensor (Invitrogen, Thermo Fisher Scientific), offer a rapid, high-throughput alternative to FLP incubations, facilitating comparisons across datasets and ecosystems. These dyes accumulate in digestive cellular compartments, enabling microscopic or flow cytometric quantification of acidic vacuole presence (Rose et al., 2004). Using fluorometry, these dyes enable simultaneous detection of chlorophyll and acidic vacuoles within individual cells, indicating potential mixotrophy. Several studies have investigated LysoTracker use for tracking mixotrophy in culture experiments and have found low positive staining in a known photoautotroph (Costa et al., 2022) or in mixotrophic organisms grown under photoautotrophic conditions (Anderson et al., 2017), supporting the application of acidotropic dyes to track active mixotrophy. However, these dyes have also been used in studies unrelated to mixotrophy, and have been shown to demonstrate non-grazing specific staining (Wilken et al., 2019). For example, LysoTracker and LysoSensor dyes have been used to selectively label acidic sites of active frustule formation in diatoms (Desclés et al., 2008; McNair et al., 2018; Shimizu et al., 2001). Additional studies using a diverse range of cultures are thus needed to better evaluate the functionality, strengths, and limitations of acidotropic dyes for detecting mixotrophs in natural marine communities.

Despite limitations, preliminary studies using LysoTracker have seemingly provided realistic insights into mixotrophic presence within the natural environment. In the North Pacific, higher LysoTracker staining (indicating higher mixotrophic presence) correlated with low-nutrient conditions and demonstrated consistent trends with FLP incubation observations from the same region (Sato et al., 2017; Sato & Hashihama, 2019). However, unexpected staining at stations with high diatom abundance hinted at the possibility of community-specific staining artifacts (Sato & Hashihama, 2019). A recent study using LysoTracker to quantify mixotroph abundance found mixotrophs correlated with prey concentration, temperature, and cell size (Ewton, 2025), consistent with described impacts of environmental factors on mixotrophs based on previous experimental findings and model predictions (Kang et al., 2020; Moeller et al., 2024; Ward & Follows, 2016; Wilken et al., 2013). Another study that combined fluorescence-activated cell sorting with FLP incubations and LysoSensor Blue staining revealed that while both methods could identify mixotrophs, LysoSensor exhibited a greater specificity and lower rate of false positives compared to FLP incubations (Florenza et al., 2024). Finally, a recent study in the Indian Ocean found mixotroph abundance, as determined with LysoTracker, correlated positively with heterotrophic bacterial prey and comprised nearly 80% of eukaryotic biomass (Selph et al., 2025). These findings highlight the potential of acidotropic dyes as ecological tools, while also underscoring the need for rigorous validation across taxa, among environments, and between acidotropic dye types.

Given the relative ease and scalability of acidotropic dye methods, our study aimed to evaluate their specificity and reliability for detecting mixotrophs *in situ.* To address this need, we tested LysoTracker Green (DND-26) and LysoSensor Blue (DND-167) across a range of photoautotrophic and mixotrophic cultures. We further tested these dyes in natural environments by comparing their staining patterns with fluorescently labeled particle (FLP) incubations across two ecologically distinct regions: the California Current System (CCS), characterized by seasonal upwelling and diatom dominance (Abdala et al., 2022; Lassiter et al., 2006), and the seasonally stratified North East Shelf (NES), which supports a more diverse protistan community (Marrec et al., 2021). Additionally, we incubated CCS seawater to determine how simulated blooms might play a role in dye staining behavior. Our findings provide new insights into the utility and limitations of acidotropic dyes for detecting mixotrophs in the field and will be valuable in guiding future field efforts.

## Methods

### Acidotropic dye culture tests

Eighteen isolates of marine phytoplankton (Table 1) were maintained in semi-continuous batch culture in nutrient replete conditions. Cultures were grown in filtered and chelexed natural Gulf Stream seawater containing Aquil* nutrient additions (Price et al., 1989; Sunda et al., 2005). Trace metals (selenium, cobalt, zinc, copper, manganese, and iron) were added at Aquil* concentrations (Sunda et al., 2005), with additional supplementation of cadmium (1.0 nM) and nickel (100 nM). All metals were buffered with 100 µM EDTA. Molybdenum was not added as it is naturally occurring in seawater (Pakalns et al., 1978; Smedley & Kinniburgh, 2017). Synthetic Aquil* media was made for *P. tricornutum* with nutrient concentrations as above using ultrapure Milli-Q base water (Milli-Q IQ 7000 and IQ Element) and Aquil* salt concentrations (Price et al., 1989), but without silica added to investigate acidotropic dye staining in a non-silica biomineralizing diatom. Sterility of Gulf Stream water was achieved through 0.2 µm filtration and microwaving of the seawater for 15 minutes before media preparation. Cultures were not treated with antibiotics, and were therefore non-axenic. Cultures were grown in various temperature and light conditions detailed in Table 1.

**Table 1:** Phytoplankton and mixotroph cultures included in this study along with their isolation locations, growth conditions, trophic mode, lineage, and approximate size range determined via flow cytometry. Sources for mixotrophic metabolism, acquisition, and size dimensions (µm) are provided for those that were available in the table footnotes. *Micromonas polaris* was too small to measure size with the Attune CytPix flow cytometer, so its dimensions are reported from published references. Organisms larger than the image field (>74 µm) are indicated with length “>74”.

| Culture | Isolation Location | Temperature (°C) | Light Cycle | Light Intensity (µmol photons m<sup>-2</sup> s<sup>-1</sup>) | Metabolism | Lineage | Acquisition | CytPix Size (Width x Length) (µm) |
| --- | --- | --- | --- | --- | --- | --- | --- | --- |
| Chaetoceros sp. 02 (UNC2302) | California Current System | 12 | 12 light: 12 dark | 150 | Phototroph | Diatom | Adrian Marchetti (UNC Chapel Hill) | 7 x 13 |
| Chaetoceros sp. 12 (UNC2312) | California Current System | 12 | 12 light: 12 dark | 150 | Phototroph | Diatom | Adrian Marchetti (UNC Chapel Hill) | 17 x >74 |
| Chaetoceros sp. 22 (UNC2322) | California Current System | 12 | 12 light: 12 dark | 150 | Phototroph | Diatom | Adrian Marchetti (UNC Chapel Hill) | 14 x 37 |
| Odontella sp. (UNC2314) | California Current System | 12 | 12 light: 12 dark | 150 | Phototroph | Diatom | Adrian Marchetti (UNC Chapel Hill) | 16 x >74 |
| Chlamydomonas sp. (CCMP235) | Nantucket Sound, Massachusetts | 12 | 12 light: 12 dark | 150 | Phototroph | Chlorophyte | Bigelow NCMA | 8 x 14 |
| Gephyrocapsa oceanica (UGA06) | South Atlantic Bight | 18 | 13 light: 11 dark | 110 | Mixotroph?<sup>1</sup> | Coccolithophore | Lucy Quirk (UGA)<sup>5</sup> | 8 x 10 |
| Gephyrocapsa huxleyi (UGA13) | South Atlantic Bight | 18 | 13 light: 11 dark | 110 | Mixotroph<sup>1</sup> | Coccolithophore | Lucy Quirk (UGA)<sup>5</sup> | 6 x 7 |
| Tetraselmis sp. | Waquoit Bay, Massachusetts | 18 | 13 light: 11 dark | 110 | Mixotroph<sup>2</sup> | Chlorophyte | Nicole Millette (VIMS) | 8 x 15 |
| Tetraselmis chui (PLY429) | Puerto Rico | 18 | 13 light: 11 dark | 110 | Mixotroph<sup>2</sup> | Chlorophyte | Millford Strain Collection | 9 x 17 |
| Odontella rostrata (UGA01) | South Atlantic Bight | 18 | 13 light: 11 dark | 110 | Phototroph | Diatom | Lucy Quirk (UGA)<sup>5</sup> | 13 x >74 |
| Phaeodactylum tricornutum (CCMP2561) | North Atlantic | 18 | 13 light: 11 dark | 110 | Phototroph | Diatom | Bigelow NCMA | 4 x 20 |
| Micromonas polaris (CCMP2099) | Baffin Bay, Arctic | 4 | 24 light | 70 | Phototroph?<sup>3</sup> | Prasinophyte | Bigelow NCMA | 1 x 3<sup>7</sup> |
| Chaetoceros neogracile (RS19) | Antarctica | 4 | 24 light | 70 | Phototroph | Diatom | Mak Saito (WHOI)<sup>6</sup> | 6 x 9 |
| Fragilariopsis cylindrus (UNC1902) | Palmer LTER, Antarctica | 4 | 24 light | 70 | Phototroph | Diatom | Adrian Marchetti (UNC Chapel Hill) | 5 x 10 |
| Pseudo-nitzschia subcurvata (UNC1901) | Palmer LTER, Antarctica | 4 | 24 light | 70 | Phototroph | Diatom | Adrian Marchetti (UNC Chapel Hill) | 7 x 31 |
| Geminigera cryophila (CCMP2564) | Ross Sea, Antarctica | 4 | 24 light | 70 | Mixotroph<sup>4</sup> | Cryptophyte | Rebecca Gast (WHOI)<sup>4</sup> | 12 x 15 |
| Mantoniella antarctica (SL-175) | Ross Sea, Antarctica | 4 | 24 light | 70 | Mixotroph<sup>4</sup> | Prasinophyte | Rebecca Gast (WHOI)<sup>4</sup> | 6 x 8 |
| Pyramimonas tychotreta (I-9 Pyram) | Ross Sea, Antarctica | 4 | 24 light | 70 | Mixotroph<sup>4</sup> | Prasinophyte | Rebecca Gast (WHOI)<sup>4</sup> | 10 x 17 |

<sup>1</sup>Ye et al. (2024), Biology <sup>2</sup>This study <sup>3</sup>McKie-Krisberg & Sanders (2014), ISME; Wilken et al. (2019), Phil Trans R Soc B; Jimenez et al. (2021), J Phycol <sup>4</sup>Gast et al. (2014), FEMS Microbiol Ecol <sup>5</sup>Quirk et al. (2025), Limnology and Oceanography <sup>6</sup>Kellogg et al. (2022), Limnology and Oceanography <sup>7</sup>Simon et al. (2017), Protist

Cell growth was monitored at least 3 times a week using flow cytometry, and cultures were stained with both acidotropic dyes when in the exponential growth phase. LysoTracker^TM^ Green DND-26 (Invitrogen, Thermo Fisher Scientific) was incubated with cultures at concentrations of 75 nM (Rose et al., 2004), and LysoSensor^TM^ Blue DND-167 (Invitrogen, Thermo Fisher Scientific) at 1 µM (Carvalho & Granéli, 2006). Both stains were incubated with cells for 10 minutes in the dark prior to measurements, with measurements performed no more than 30 minutes after initial stain introduction. An additional test of the staining of cultures in stationary, exponential, and after 24 hours of darkness was performed in the same manner (Supplemental Figure 1). Flow cytometric measurements were primarily conducted using an Attune CytPix Acoustic Focusing and Imaging Cytometer (Thermo Fisher Scientific) equipped with violet (405 nm), yellow (561 nm), and blue (488 nm) lasers. Thirteen of nineteen cultures were also analyzed on a Guava easyCyte 5HT (Cytek Biosciences; blue 488 nm and green 532 nm lasers) to allow for comparisons with field data, which were collected on this instrument, and to determine if staining varied by instrument type. As LysoSensor Blue requires violet excitation, these measurements were only possible on the Attune CytPix. Cell size was estimated on the CytPix using the “Cells Full Resolution (v34)” model, which may slightly overestimate cell sizes (Software v7.1), but were determined to be comparable to a Coulter Counter (data not shown). Reported values demonstrate the minor to mean major diameter as a proxy for cell width and length, respectively (Table 1). Cells larger than the 74 µm image field are indicated as >74 µm. *Micromonas polaris* was too small for size determination on this instrument. On the Guava easyCyte, cells larger than ∼20 µm were not captured.

All cells were identified as stained or unstained based on gates that were similar to those applied in the field (Supplemental Figure 2). However, three large (>50 µm) diatoms (*Chaetoceros sp. 02, Odontella sp.,* and *O. rostrata*) were already past the fluorescence gate threshold before stain application, necessitating a higher gating threshold for both dyes. This could be due to more green autofluorescence in large cells and larger vacuoles for stain to accumulate in (Tang & Dobbs, 2007) (Supplemental Figure 2). Additionally, only one picoeukaryote was included in this study, *Micromonas polaris*, and was given a picoeukaryote specific gate as smaller cells naturally have less green autofluorescence and smaller vacuoles (Supplemental Figure 2). The three large diatoms were measured exclusively on the CytPix as it can better accommodate larger cells (Luminex Corporation, 2017; Scientific, 2022).

The number of biologically replicated runs performed with LysoTracker and LysoSensor ranged between 2 and 9, dependent on growth of the organism over the testing period (Supplemental Table 1). Between 100 and 700,000 (median 6,000) cells were assessed for staining within each biological replicate (Supplemental Table 1). Each biological replicate was run in stained technical duplicates and positively stained cells were identified based on high red fluorescence, and high green fluorescence for LysoTracker or high blue fluorescence for LysoSensor (Supplemental Table 2). Controls of unstained samples were also run in technical duplicates and used to correct for natural fluorescence of the cells. The percent of positively stained chlorophyll-containing cells (*P*) was calculated as the percent of cells in the stained region (high green or blue fluorescence) in the stained sample minus that in the unstained control.

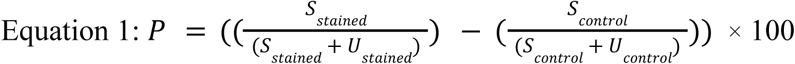

Where *S* is the concentration of chlorophyll-containing cells within the dye-stained region and *U* is the concentration of these cells in the unstained region, for both stained and control samples. The final reported percent of stained cells is calculated as the average and standard deviation of biological replicates.

A quantification of the degree of staining of each culture was also assessed by taking the difference between mean green or blue fluorescence in control samples from those in stained samples. The reported values are the average of two technical replicates. Due to some changes in fluorescence being zero, a small integer (0.001) was added to all values to allow plotting on a log scale.

### Prey removal assays

Grazing assays were performed with three known mixotrophs from the Southern Ocean (*G. cryophila, M. antarctica,* and *P. tychotreta*) and two organisms from the *Tetraselmis* genus (Supplemental Figures 3-4) using the established prey removal framework (Frost, 1972). These grazing assays used two treatments: prey alone and prey with mixotrophic predators, where changes in prey growth rates in the presence and absence of predators serve as an indirect measurement of predator ingestion rates. These experiments were conducted with cells grown into an early exponential phase (350 mL for Southern Ocean mixotrophs and 1 L for *Tetraselmis spp.*). At the start of the grazing experiments, a portion of the volume was removed and filtered through a 0.7 µm GF/F filter and returned to the batch culture to serve as a source of dissolved organic matter (DOM) (100 mL for Southern Ocean mixotrophs and 250 mL for *Tetraselmis spp*). DOM was added to promote bacterial growth, as the bacteria co-cultured with the phytoplankton rely on organic substrates derived from these cells. Triplicate flasks were then prepared for the mixotroph and prey treatment (50 mL for Southern Ocean mixotrophs and 100 mL for *Tetraselmis spp*). The remaining culture volume was filtered to remove mixotrophs using a 3 µm (*P. tychotreta* and *G. cryophila*) or 1 µm (*M. antarctica* and *Tetraselmis spp.*) polycarbonate filter. The filtered volume, now devoid of mixotrophs, was used to prepare triplicate flasks for the prey only treatment. For the Southern Ocean mixotrophs, experimental timepoints were collected at 24 and 48 hours due to slower acclimation periods at cold temperatures. For *Tetraselmis spp.,* sampling timepoints were taken at 0 and 24 hours. Approximately 3 mL triplicate technical replicates were collected and preserved with 0.25% glutaraldehyde for the Southern Ocean mixotroph samples and 0.5% buffered paraformaldehyde for the *Tetraselmis spp.* samples. Mixotroph and bacterial abundances were quantified using a BD C6 Accuri flow cytometer (BD Biosciences). For calculations, the average mixotroph concentration was used between the two timepoints as it was within error compared to equations taking predator growth into account (Heinbokel, 1978). Prior to analysis, bacteria were stained with 1x SyBr Green (Fisher). Ingestion rates were expressed as bacteria consumed per mixotroph per hour (Frost, 1972).

To further confirm prey ingestion in *Tetraselmis*, which has not previously been reported, FLP incubations were performed on the two *Tetraselmis spp.* isolates. Fluorescent microspheres (0.5µm diameter, Fluoresbrite, Polysciences) were added at a concentration of 500,000 particles mL^−1^ into 50 mL of culture. Immediately after the spike of microspheres, triplicate samples were taken for a T_initial_ timepoint and preserved using 4% ice cold glutaraldehyde. Cultures were then incubated with the microspheres for an hour, sampled again for T_final_, and preserved in the same manner. Preserved samples were filtered onto 25 mm 3 µm polycarbonate filters, mounted to microscope slides using DAPI Vectashield (ThermoFisher), and stored at -20°C. Slides were counted using a Nikon Eclipse Ti2 microscope with NIS element AR 6.10.01 on the FITC (466 nm excitation, 520 nm emission) and TRITC (561 nm excitation, 600 nm emission) channels. Fifteen fields of view at 1.7mm^2^ each were imaged and examined, resulting in 150-750 cells examined for bead ingestion. Percent of the population that was mixotrophic was calculated by dividing the total number of examined cells that had ingested a bead by the total number of cells examined. For each biological replicate, the T_final_ value was subtracted from the T_initial_ value. A z-stack image was captured for *Tetraselmis sp.* containing a fluorescent bead (Supplemental Figure 4) with the same channels as above, with 20 slices collected over a 10 µm range.

### Measurements at sea

The North East Shelf (NES) was sampled along the Long Term Ecological Research (LTER) transect during July 13th - July 18th, 2022 aboard the *R/V Endeavor* cruise EN685 (Figure 1a). Seawater was collected at the surface (3 meters) and the subsurface chlorophyll maximum (SCM, 7-27 meters). At three stations (L06, L10, LX), two additional depths were also sampled to achieve more depth resolution. These depths were below the SCM and ranged from 30-45 meters.

**Figure 1:**
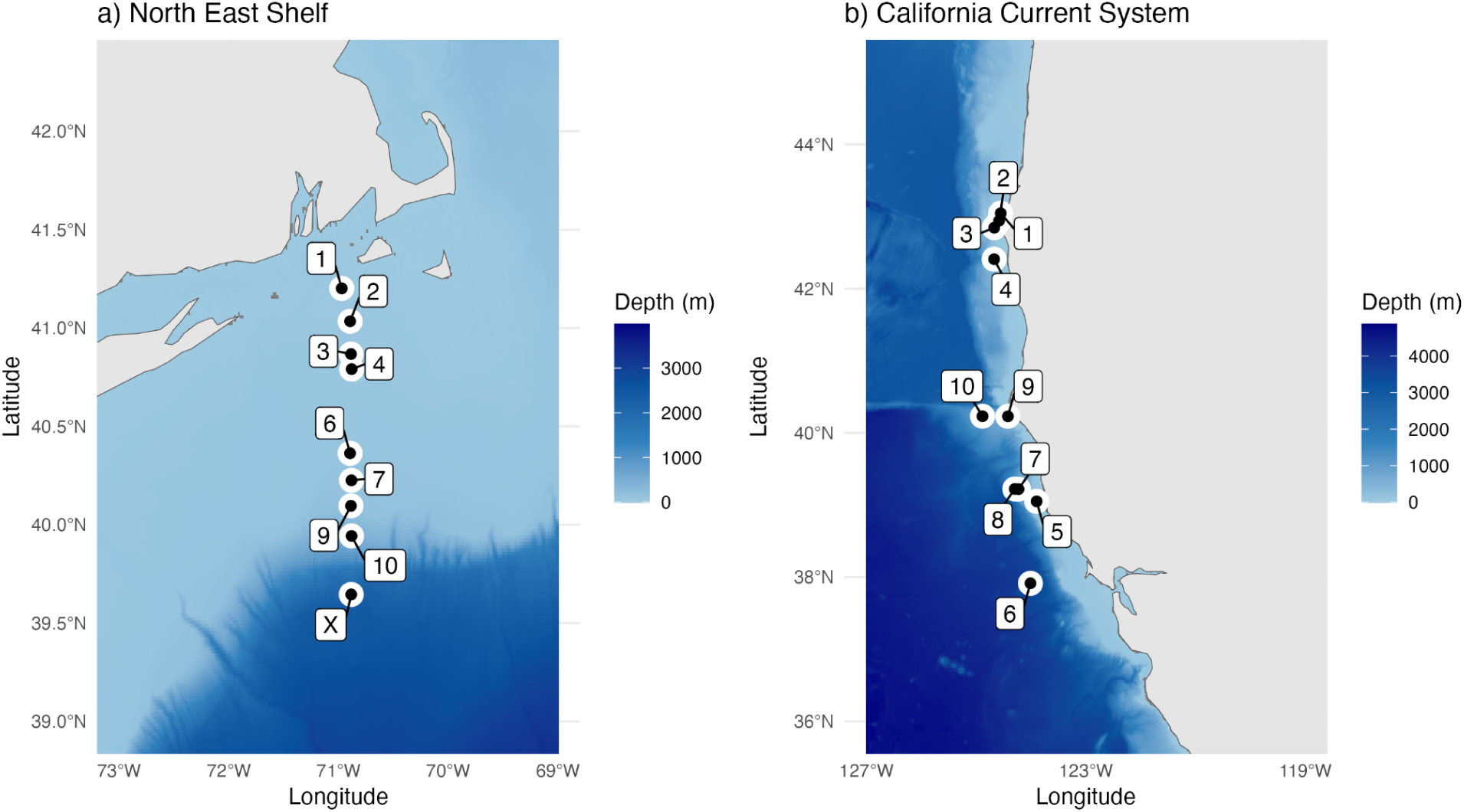
Sampling locations for both the North East Shelf (a) and California Current System (b) cruises. Color represents depth in meters, with individual scales for both plots.

The California Current System (CCS) was sampled aboard the *R/V Sally Ride* as part of the Phytoplankton response to the UPwelling CYCLE (PUPCYCLE II) cruise SR2311 from May 28th - June 11th, 2023 (Figure 1b). The goal of the CCS expedition was to sample upwelled water and follow upwelled filaments as they aged (Lin et al., 2024; Till et al., 2019, 2025). Samples were collected at 46%, 22%, 10% and 1% of surface irradiance (7-69 meters).

LysoTracker data from both the NES and CCS have previously been reported using a similar dataset focused on the subsurface only for the CCS (10% surface irradiance) (Ewton, 2025). Here, we build on this by comparing direct grazing estimates (FLP incubations) to LysoTracker results.

Mixotroph presence was determined using LysoTracker Green. The dye was applied to live seawater samples at a final concentration of 75 nM in the same manner as the culture experiments (Rose et al., 2004). For NES samples, seawater was filtered through a 200 µm mesh before LysoTracker addition, and through a 52 µm mesh directly before flow cytometry injection to avoid clogging the flow cell. For CCS samples, water was pre-filtered through 52 µm mesh before staining with LysoTracker. Stations 1 through 4 on the CCS cruise do not have LysoTracker derived data.

Flow cytometry was used to analyze LysoTracker stained seawater on both cruises, though slightly different settings and Guava easyCyte HT flow cytometers were used (Supplemental Table 2). Communities of interest were gated based on community recommendations (Thyssen et al., 2022) (Supplemental Figure 5). Briefly, *Synechococcos* were detected based on high yellow fluorescence and low forward scatter (FSC), picoeukaryotes (∼1-3 µm) were detected based on medium yellow and red fluorescence, and nanoeukaryotes (∼3-20 µm) were detected based on high red fluorescence (Supplemental Figure 5). Mixotroph identification was detected based on high red and high green fluorescence when LysoTracker stain was applied.

Heterotrophic nanoeukaryotes were identified based on low red and high green fluorescence when LysoTracker was applied (Rose et al., 2004). A threshold for event collection was set to a very low red fluorescence intensity, which likely excluded small heterotrophic nanoeukaryotes that lack chlorophyll. However, heterotrophs may still register low levels of red fluorescence on the Guava EasyCyte due to signal overlap between fluorescence channels (Perfetto et al., 2012). For example, pigments that are excited by the blue laser can occasionally be weakly detected in the red channel, even in the absence of chlorophyll (Luminex Corporation, 2017; Macel et al., 2020). As a result, the heterotrophic population reported here includes primarily larger cells with spectral overlap or red fluorescence not related to chlorophyll that was strong enough to exceed the red detection threshold.

### Fluorescently labeled particle incubations at sea

FLP experiments were conducted on the NES cruise using seawater from two depths (surface and SCM), and on the CCS cruise at one depth (surface). For each incubation, 500 mL of seawater was pre-screened through a 200 µm mesh to remove mesozooplankton. Fluorescent surrogate prey particles were spiked into this volume at concentrations of 10^5^ particles mL^−1^. Fluorescent microspheres (0.5 µm diameter, Fluoresbrite, Polysciences) were primarily used, however, fluorescent *E. coli* (1 µm diameter, K-12 strain, Invitrogen) was also used at all stations on the NES and demonstrated similar trends to the microspheres (Supplemental Figure 6). Immediately after prey addition, a time-zero (T_initial_) sample was taken and preserved using 4% glutaraldehyde. Incubations were then carried out for one hour in mesh bags in a clear plexiglass incubator with flow-through seawater to maintain *in situ* temperature and light availability. After an hour incubation another sample (T_final_) was taken and preserved in the same manner as T_initial_.

On the NES cruise, 2 mL subsamples from T_initial_ and T_final_ were analyzed on a Guava EasyCyte HT flow cytometer to quantify photosynthetic cells co-associated with a green fluorescent particle tracer (Supplemental Figure 7). Instrument gain settings are listed in Supplemental Table 2. Preserved samples from stations L1 and L2 on the NES cruise and stations 9 and 10 on the CCS cruise were also run on an Attune CytPix to gather qualitative photos of the large nanoeukaryotes. Photos demonstrate organisms with high red fluorescence and FSC. A random subset of 49 photos (each 74 x 74 µm) was collected to illustrate differences in community composition between cruises (Figure 5 e, f).

For the CCS cruise, microscopy was used for analyzing FLP ingestion. Volumes of 10-30 mL were fixed, filtered onto 3 µm polycarbonate filters, and mounted using DAPI Vectashield (Vector Laboratories). Epifluorescent microscopy was used to manually determine mixotrophic cells, those that contained chloroplasts and green fluorescent prey, as well as photoautotrophic cells that had only chloroplasts (no intracellular prey observed). Samples were enumerated using a field-of-view counting method with a calibrated reticle on an Olympus CKX53 inverted microscope at 400x total magnification. For each sample, a minimum of 300 chloroplast-containing cells were enumerated to ensure statistical reliability.

Although NES FLP analyses were primarily flow cytometric, microscopy slides were also prepared to enable comparisons across cruises (Supplemental Table 3). Microscopy slides were collected in the same manner as on the CCS. These slides used *E. coli* prey, which exhibited clumping due to freezing and thawing, and were more challenging to interpret. NES slides were analyzed using an Olympus BX60 compound microscope at 1000x total magnification, with cell counts performed along one to three transects across the central portion of the slide. The transect method proved unreliable for generating cell concentrations as it required manual stage movements by minute increments, which made the effective counting area difficult to reproduce accurately. As a result, only the proportion of mixotrophic cells will be used from NES FLP microscopy samples. These different counting methods and microscopes from the CCS were due to using a newer microscope model not available at the time of the NES evaluations.

From this data, we calculated the percent of mixotrophic phototrophs, mixotroph abundance, cell specific grazing rates, and community bacterivory rates. Mixotroph abundance was calculated by the Guava EasyCyte HT for NES flow cytometry samples, and was manually calculated for microscopy samples as follows:

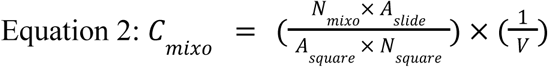

where C_Mixo_ is the concentration of mixotrophs, N_mixo_ is the number of mixotrophs counted, A_slide_ is the total filtration area, A_square_ is the area of the square being counted at the objective used, N_square_ is the total number of squares counted, and V is the volume of water filtered. All reported mixotroph values (percent and abundance) were calculated by subtracting the T_initial_ samples from those at T_final_. Percent of mixotrophic phototrophs was calculated by dividing the mixotrophic cells by all autotrophic cells and subtracting the T_initial_ value from the T_final_.

Reported is the cell specific grazing rate (CSGR) out of all photosynthetic organisms (Koppelle et al., 2024; Sherr et al., 1988; Thurman et al., 2010).

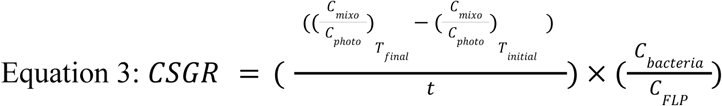

Where the concentration of mixotrophs (C_mixo_) divided by the total number of pigmented phototrophs (C_photo_) for the initial (T_initial_) timepoint was subtracted from the final timepoint (T_final_), and divided by the total time (t) of the incubation in hours. The bacteria to prey ratios were then multiplied to estimate the concentrations of bacteria consumed by mixotrophs where C_bacteria_ is the natural concentration of heterotrophic bacteria and C_FLP_ is the concentration of fluorescently labeled particles spiked into the sample. An ingestion rate of 1 prey per mixotroph was assumed to prevent biases between analysis methods, as flow cytometry cannot determine how many prey particles were ingested per mixotrophic cell. Cell specific grazing rates are reported in bacteria mixotroph^−1^ hour^−1^. Natural bacteria concentrations were acquired using additional samples taken on each cruise and preserved with 0.25% glutaraldehyde, frozen at -80° C, and stained with 1x SyBr Green (Fisher), and measured on the Guava EasyCyte HT flow cytometer.

As another grazing metric, community bacterivory rate (*BR*), was also assessed out of all photosynthetic organisms (Li et al., 2023; Princiotta & Sanders, 2017; Unrein et al., 2014).

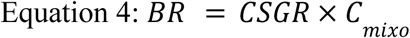

Where the community bacterivory was calculated by multiplying CSGR by the total concentration of mixotrophs (C_mixo_). For C_mixo_, the background concentration at T_initial_ was subtracted from the measured concentration at T_final_. Community BR is reported in units of bacteria mL^−1^ hour^−1^. This metric may also be referred to as grazing impact (Unrein et al., 2007, 2014).

### CCS Incubation Experiment

During the CCS cruise, incubation of actively upwelling water was performed at station 2 to simulate a natural bloom (Figure 1). This water was pumped from below the euphotic zone at 55 m depth and used to fill 20 L cubitainers as part of a larger study (Speciale, 2025). Biological triplicate cubitainers were sampled for nanoeukaryote abundance and nitrate concentrations at days 0, 2, 7, and 11. LysoTracker staining was performed at timepoints 7 and 11 in the same manner as along the CCS transect. FLP was also performed in the same manner as along the CCS transect, however, there was extremely low grazing on fluorescent beads and grazing values were unable to be reliably quantified.

Metatranscriptome analyses were performed in the CCS both within the incubation experiment and along the transect to determine community composition (Speciale, 2025). Surface niskins (46% surface irradiance) were subsampled during daily casts (Figure 1) along with each timepoint of the incubation experiment for RNA extraction. For this, 2.5 L to 4 L of seawater were filtered onto 142 mm 0.8 µm Pall Supor filters using a peristaltic pump, flash frozen, and stored at -80 °C until extraction. RNA was extracted using RNAqueous-4PCR kit and sequencing was performed on an Illumina HiSeq 4000 with 2×150 bp configuration (Speciale, 2025). Briefly, raw reads were trimmed, individual assemblies were created using rnaSPAdes, an assembly was merged and combined with CD-HIT, and annotations were performed with the Marine Functional Eukaryotic Reference Taxa (MarFERRet) database (Groussman et al., 2023). The mean raw reads that were mapped and annotated can be found in Supplemental Figure 8. Contigs with no taxonomic annotation or mapped to Opisthokonta were removed prior to downstream analysis. DESeq2 was used to obtain normalized counts for species and lineages (Love et al., 2014). Full bioinformatic details are reported in Speciale (2025).

### Statistical analysis

All data analysis and visualization were performed in R (R Version 4.3.1; R Studio Version 2024.12.1.563). Analyses utilized several packages including dplyr v1.1.4 (Wickham et al., 2023), tidyr v1.3.1 (Wickham et al., 2024), ggplot2 v3.5.2 (Wickham, 2016), gt v1.0.0 (Iannone et al., 2025), RColorBrewer v1.1-3 (Neuwirth, 2022), patchwork v1.3.0 (Pedersen, 2024), gtools v3.9.5 (Warnes et al., 2023), ggpubr v0.6.0 (Kassambara, 2023), ggpmisc v0.6.1 (Aphalo, 2024), and gridExtra v2.3 (Auguie, 2017). To determine statistical differences between cruise datasets, normality was first assessed using Shapiro-Wilk tests. If data were normal, a two-sample t-test was applied. If data were not normal, a Wilcoxon Rank-sum test was run. Bland-Altman analyses were performed with mean bias, standard deviation of differences, and 95% limits of agreement (LoA) calculated. A p-value less than or equal to 0.05 was considered statistically significant.

## Assessment

### Acidotropic dye culture experiments

To assess phagocytic potential across a diverse set of cultured marine phytoplankton, LysoTracker and LysoSensor acidotropic dyes were tested in a set of 18 phytoplankton cultures, representing five major taxonomic groups: chlorophyte (*n*=3), coccolithophore (haptophyte) (*n*=2), cryptophyte (*n*=1), diatom (*n*=10), and prasinophyte (*n*=3).

We utilized five species we knew to be mixotrophic, three of which were shown to consume prey in a previous study (*Geminigera cryophila, Mantoniella antarctica, Pyramimonas tychotreta*; McKie-Krisberg et al., 2015) and were confirmed to ingest prey in this study (Supplemental Figure 3). Two species of the *Tetraselmis* genus were further demonstrated to consume prey in this study (Supplemental Figure 4). These known mixotrophs showed consistently high LysoTracker staining, with 87–95% of cells staining positively (Figure 2a; Supplemental Table 1). Further, we tested *Gephyrocapsa huxleyi* and *Gephyrocapsa oceanica*, which were assumed to be mixotrophic due to ingestion of FLP by similar species in previous literature (Godrijan et al., 2020; Koester et al., 2025; Rokitta et al., 2011; Ye et al., 2024). These two coccolithophores demonstrated high staining between 70 and 94% (Figure 2a; Supplemental Table 1). In contrast, LysoSensor stained fewer mixotrophs overall, with only *G. cryophila* showing strong positive staining (∼93%), and *Tetraselmis spp.* staining moderately (49–69%) (Figure 2b; Supplemental Table 1). Other mixotrophic taxa (*G. oceanica, G. huxleyi, P. tychotreta, M. antarctica*) showed little or no LysoSensor signal (0-15%). These results suggest that LysoTracker more accurately stains acidic vacuoles associated with phagotrophy in mixotrophs.

**Figure 2:**
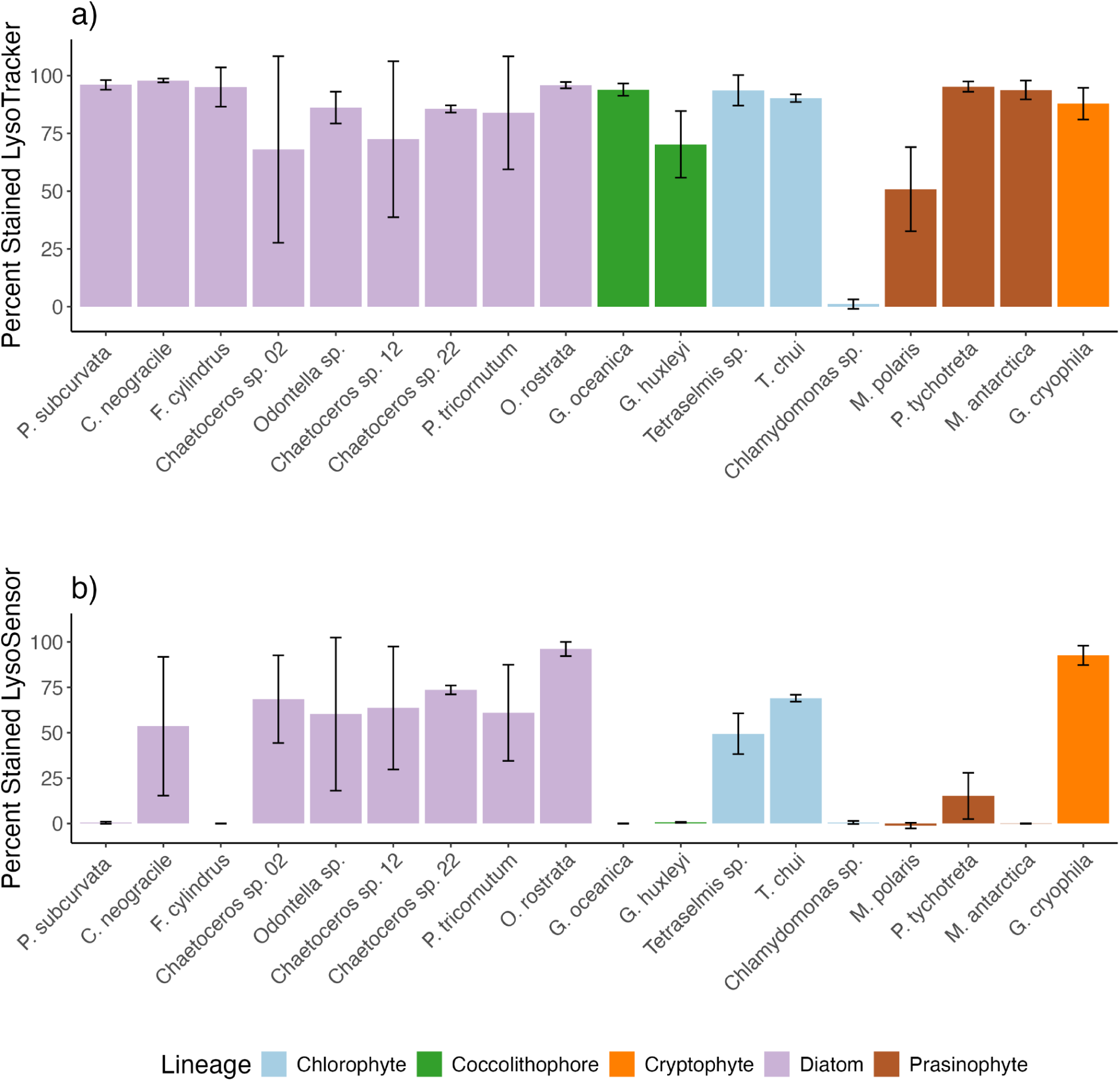
Percent of cells positively stained with LysoTracker (combined average of Guava and CytPix values) (a) or LysoSensor (b). Bars represent the average of all biological replicates and error bars indicate the standard deviation of biological replicates. The color of the bar indicates lineage. Note that gating thresholds for both dyes differed among large diatoms, nano-sized eukaryotes, and the picoeukaryote *M. polaris* (Table 1, see methods).

We also evaluated species assumed to be strictly photoautotrophic. Two non-diatom species, *Micromonas polaris* (prasinophyte) and *Chlamydomonas sp.* (chlorophyte), were assumed to be strictly photoautotrophic based on previous literature (Bock et al., 2021; Costa et al., 2022; Jimenez et al., 2021; Wilken et al., 2019). *Chlamydomonas sp.* did not stain with either dye (≤1%), supporting its photoautotrophic characterization (Figure 2; Supplemental Table 1). *Micromonas polaris* demonstrated ∼50% of cells stained with LysoTracker and no staining with LysoSensor (Figure 2; Supplemental Figure 2). LysoTracker stained all ten strictly photoautotrophic diatoms tested, with many showing >90% staining. LysoSensor, by contrast, stained only a subset of diatoms, and often at lower levels (0–83%). Notably, *Fragilariopsis cylindrus* and *Pseudo-nitzia subcurvata* showed no LysoSensor signal, despite staining strongly with LysoTracker (Figure 2b; Supplemental Table 1).

Growth phase demonstrated some changes in staining dynamics in some species (Supplemental Figure 1). For example, with LysoTracker, both coccolithophores demonstrated more staining during exponential growth and *Odontella sp.* demonstrated more staining during stationary growth. LysoSensor staining was the lowest for both *Tetraselmis spp.* during the stationary phase, and decreased slightly for *G. cryophila* after 24 hours in darkness.

### Fluorescence changes induced by acidotropic dyes

To better understand staining dynamics, we examined changes in mean green fluorescence following LysoTracker addition to assess whether cultures that produced false positives exhibited unusually high or low fluorescence signals (Figure 3a). The photoautotroph *Chlamydomonas sp.* did not exceed the predetermined flow cytometry threshold for positive staining and showed minimal changes in green fluorescence (Supplemental Figure 2, Supplemental Table 4). *Micromonas sp.* demonstrated low overall changes in green fluorescence (Supplemental Table 4), but when normalized to FSC as a proxy for size, demonstrated shifts in normalized green fluorescence that were similar to that of the other small mixotrophs (*M. antarctica, G. oceanica, G. huxleyi*), highlighting the need for size-class specific gates (Supplemental Figure 2). Taxa that exhibited a lower percent of stained cells (∼50% or lower), such as *G. huxleyi*, exhibited relatively modest increases in fluorescence intensity relative to FSC following LysoTracker staining. In contrast, *M. antarctica* displayed a high percent of stained cells despite only a small fluorescence change relative to FSC, reflecting proximity to the gating threshold prior to staining.

**Figure 3:**
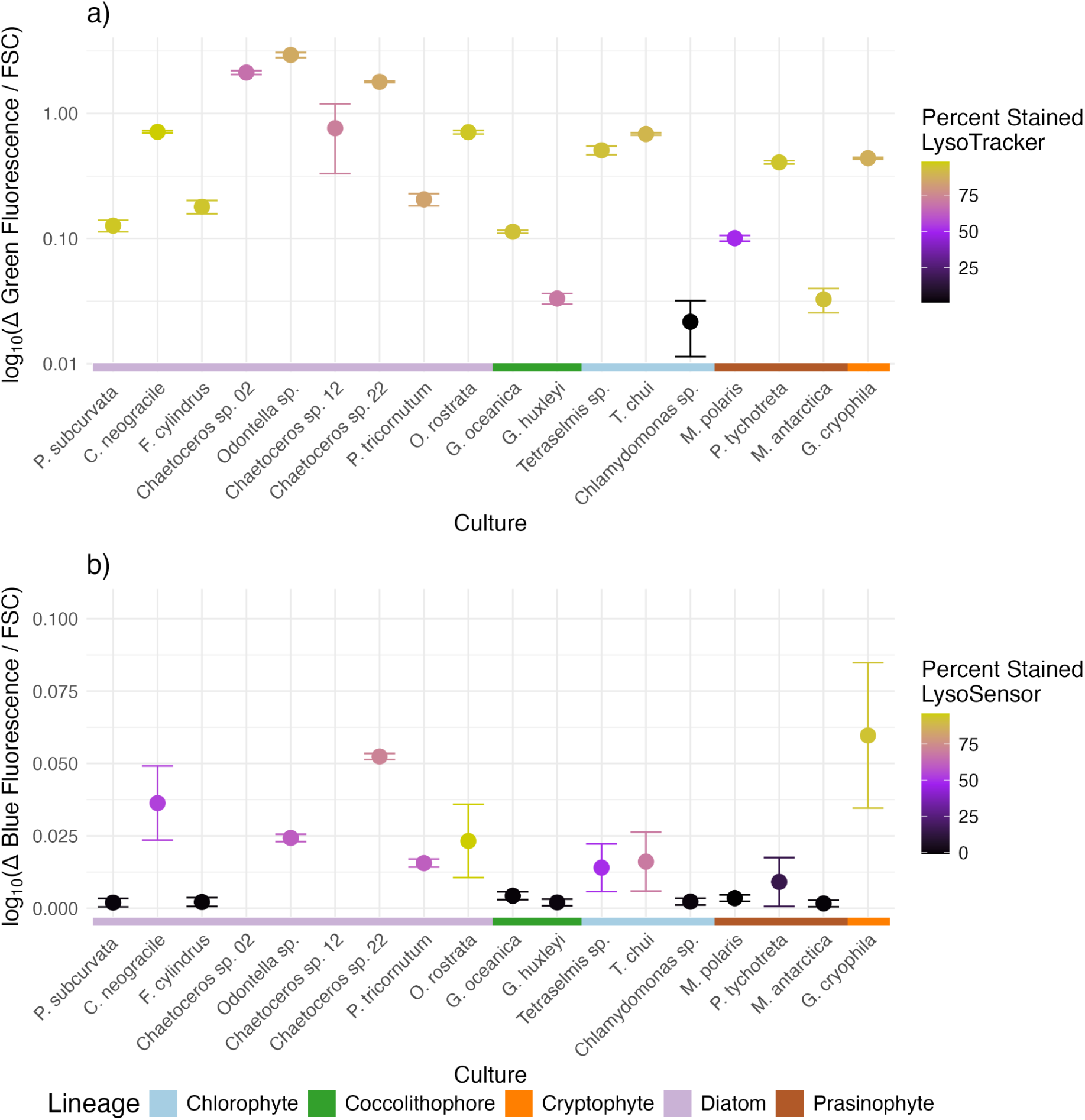
Change (Δ) in the mean green fluorescence cell^−1^ between unstained (control) and stained samples with LysoTracker (a) and change (Δ) in mean blue fluorescence cell^−1^ between unstained (control) and stained samples with LysoSensor (b) averaged between two technical replicates. Change in fluorescence for both dyes was normalized by the mean forward scatter (FSC) of each culture and plotted on a log_10_ scale. Color of dots correspond to the average percent of cells that were identified as potentially mixotrophic from all biological replicates surpassing the pre-defined gating threshold for each acidotropic dye. Note that gating thresholds for both dyes differed among large diatoms, nano-sized eukaryotes, and the picoeukaryote *M. polaris* (Table 1, see methods).

The largest increases in green fluorescence relative to FSC upon the addition of LysoTracker was observed in large chain-forming diatoms, including all tested *Chaetoceros spp.* and *Odontella spp.* All mixotrophs, except *M. antarctica*, also demonstrated relatively high changes in green fluorescence relative to FSC along with high percentages of stained cells (*Tetraselmis spp., P. tychotreta,* and *G. cryophila*).

We also examined the blue fluorescence changes following LysoSensor addition. Organisms that demonstrated no staining with LysoSensor (*P. subcurvata, F. cylindrus, G. oceanica, G. huxleyi, Chlamydomonas sp., M. polaris, P. tychotreta, M. antarctica*) also demonstrated the lowest changes in blue fluorescence relative to FSC post LysoSensor addition (Figure 3b). *P. tychotreta* demonstrated high variability, consistent with the variability seen in overall staining (Figure 2). All organisms that demonstrated staining with LysoSensor demonstrated high changes in blue fluorescence relative to FSC, with the largest fluorescent increases being in *Chaetoceros sp. 02, Chaetoceros sp. 12,* and *G. cryophila.* These results demonstrate that large chain-forming diatoms would be classified as potentially mixotrophic using either LysoTracker or LysoSensor regardless of the gating methods used.

### Cruise results

LysoTracker staining and FLP incubations were compared on the CCS and NES cruise, which builds on a previously published LysoTracker dataset from this CCS cruise and two in the NES (Ewton, 2025). The abundance of *Synechococcus*, potential prey for nanoeukaryotes, was 30-fold higher in the NES than the CCS. In the NES, *Synechococcus* values ranged 490-189,240 cells mL^−^¹ (mean: 36,090 ± 43,370), compared to 60-4,700 cells mL^−^¹ (mean: 1,200 ± 940) in the CCS (Wilcoxon p = 0.00) (Figure 4a). Similarly, heterotrophic bacteria were significantly elevated by 1.9-fold in the NES (mean: 1,307,350 ± 353,260 cells mL^−^¹) compared to CCS (mean: 698,670 ± 287,120 cells mL^−^¹) (Wilcoxon p = 0.00) (Figure 4b). Heterotrophic nanoeukaryotes were also about 6-fold higher in the NES (range: 0-2,200, mean: 700 ± 610 cells mL^−1^) than in the CCS (range: 20-540; mean: 110 ± 100 cells mL^−^¹) (Wilcoxon p = 0.00) (Figure 4c). Photosynthetic nanoeukaryote abundance did not vary significantly between cruises, with a range of 60-5,170 cells mL^−^¹ (mean: 2,000 ± 1,370) in the CCS and 340-6,290 cells mL^−^¹ (mean: 2,200 ± 1,800) in the NES (Wilcoxon p = 0.84) (Figure 4d).

**Figure 4:**
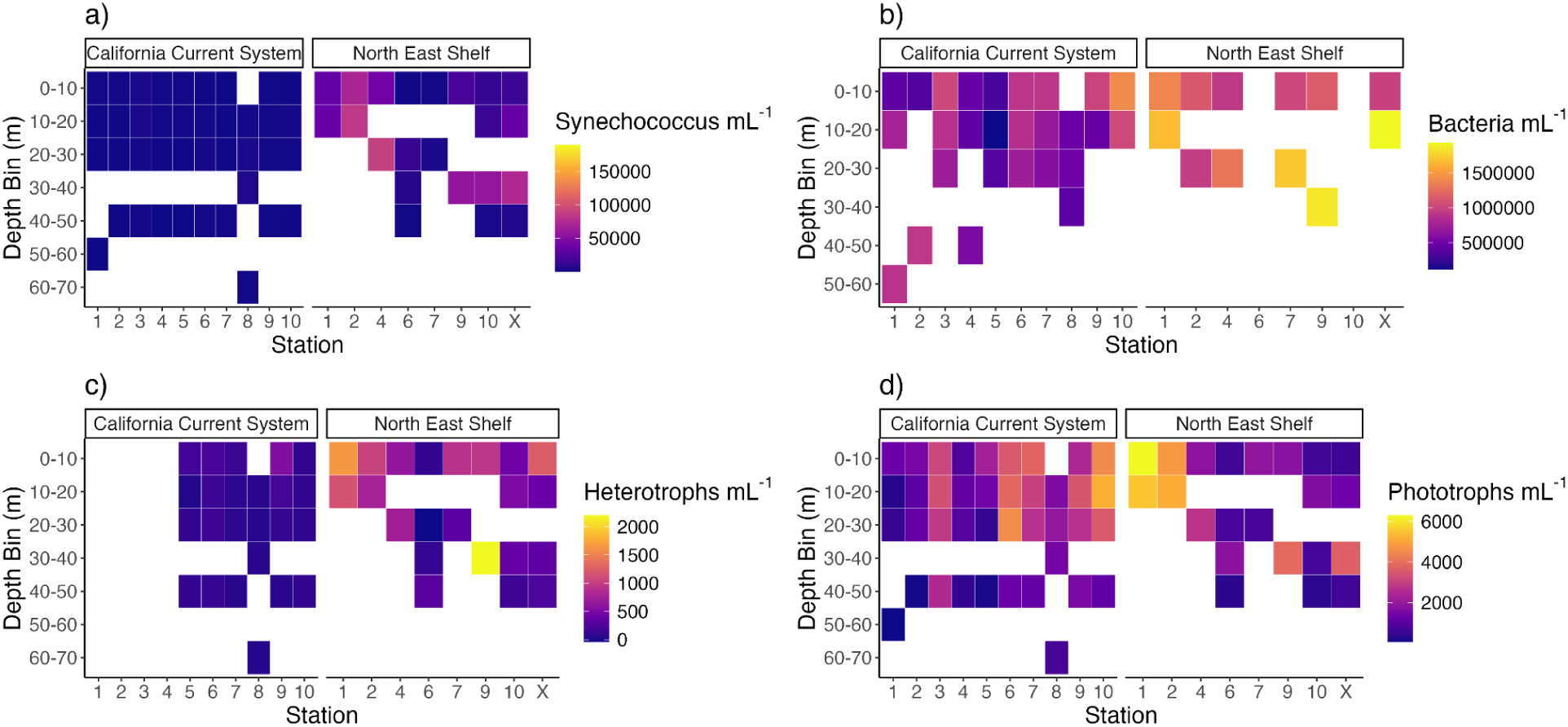
Abundance of *Synechococcus* (a), heterotrophic bacteria (b), large heterotrophic nanoeukaryotes (c), and phototrophic nanoeukaryotes (d) over the two cruise tracks. Reported values demonstrate the average value of two biological replicates. Sampled depths were aggregated over 10 m depth bins, only one sample was present per bin.

**Figure 5:**
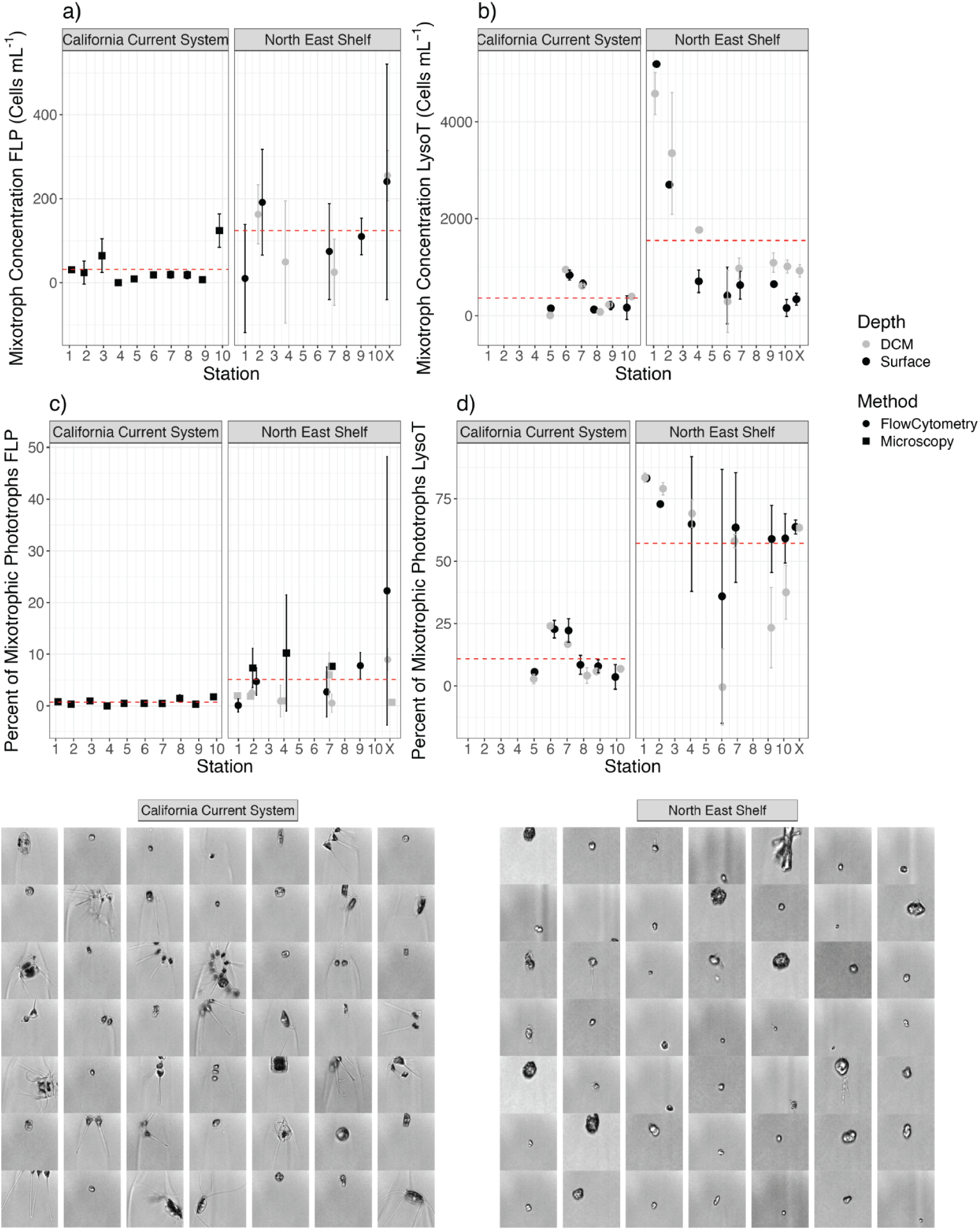
Mixotrophic abundance and percent of photosynthetic cells across the two cruise transects. Mixotroph concentration (a, b) and percent of mixotrophs within the photosynthetic community (c, d) were estimated using FLP incubations (a, c) and LysoTracker staining (b, d). Error bars represent the standard deviation of biological replicates. Colors indicate depth (gray, NES subsurface chlorophyll maximum and CCS 20% of surface light availability; black, surface). Shapes indicate FLP analysis method (circle, flow cytometry; square, microscopy). Red dashed lines indicate mean for each cruise. Bottom panels show example images (each 74 x 74 µm) of large photosynthetic nanoeukaryotes (high FSC and red fluorescence) from the Attune CytPix at CCS stations 9-10 (left) and NES station L1-L2 (right).

Mixotroph abundance was also significantly different between cruises and demonstrated significantly higher abundance, percent of the phototrophic community, and grazing in the NES compared to the CCS (Figure 5). LysoTracker stained mixotrophs were 3.5-fold higher in concentration in the NES (range: 80-4,590; mean: 970 ± 1,140 cells mL^−^¹) than in the CCS (range: 0-950; mean: 270 ± 270 cells mL^−^¹) (Wilcoxon p = 0.00) (Figure 5b). Similarly, the percent of photoautotrophic nanoeukaryotes that were considered mixotrophic with LysoTracker staining was 5-fold higher in the NES (range: 0-83.4%; mean: 48.9 ± 22.2%) than in the CCS (range: 0.0-24.0%; mean: 9.3 ± 6.8%) (Wilcoxon p = 0.00) (Figure 5d).

FLP incubation data also indicated that mixotrophic nanoeukaryotes were generally more abundant in the NES than in the CCS, despite large variability between biological replicates. In the limited NES FLP data obtained with microscopy, the mixotroph percentage was 6.5-fold higher than in the CCS, representing 4.58 ± 3.65% of the total nanoeukaryote population versus 0.72 ± 0.54% in the CCS (Wilcoxon p = 0.00) (Figure 5c). Similarly, FLP data analyzed by flow cytometry showed mixotroph concentrations in the NES (range: 10-250; mean: 120 ± 90 cells mL^−^¹) were 4-fold higher than those measured in the CCS using microscopy (range: 0-120; mean: 30 ± 40 cells mL^−^¹) (Figure 5a). Absolute mixotroph concentrations derived from microscopy on the NES cruise were not reliable (see methods). While these results point to higher FLP-derived mixotroph concentrations in the NES, the use of different analytical methods (microscopy vs. flow cytometry) between cruises means potential method-related biases cannot be ruled out. Prior work has shown that microscopy and flow cytometry estimates generally align for FLP based methods, however, these findings should be interpreted cautiously (Anderson et al., 2017; Li et al., 2023).

In addition to higher mixotrophic percentage and abundance in the NES, there were significant differences in the grazing assessed with FLP incubation methods. In the NES, cell specific grazing rate (CSGR) measured with flow cytometry was 9.5-fold higher (range: 0.01-2.19; mean: 0.67 ± 0.79 bacteria cell^−^¹ hour^−^¹) than in the CCS where microscopy was used (range: 0-0.28; mean: 0.07 ± 0.08 bacteria cell^−^¹ hour^−^¹) (Wilcoxon p = 0.01). While method-based biases may play a role, CSGR was also significantly higher in the NES than the CCS when only microscopy data was considered (Wilcoxon p = 0.01) (Supplemental Figure 9). (Note this was able to be calculated for NES microscopy data as CSGR does not take into account absolute mixotroph concentration). Additionally, CSGR was not significantly different between flow cytometry and microscopy analysis methods in the NES (Wilcoxon p = 0.74). Similarly, the population-level bacterivory rate (BR) was 36-fold greater in the NES flow cytometry data (range: 10-885; mean: 195 ± 290 bacteria mL^−^¹ hour^−^¹) compared to the CCS microscopy data (range: 0-40; mean: 5.0 ± 11.2 bacteria mL^−^¹ hour^−^¹) (Wilcoxon p = 0.00) (Supplemental Figure 10). This demonstrates that mixotrophs were more abundant and grazing more in the NES compared to the CCS by metrics measured.

Agreement between FLP and LysoTracker methods was determined using *R^2^* values and Bland-Altman tests. The highest agreement was observed for CCS percent mixotrophs (*R^2^* = 0.20), while the lowest was for CCS mixotroph concentration (*R^2^* = 0.05), with NES values intermediate (*R^2^* = 0.13 and 0.11) (Supplemental Figure 11). Across all measurements, LysoTracker produced higher concentrations and percentages than FLP. Bland–Altman analysis compared the differences between FLP and LysoTracker estimates, showing smaller discrepancies for CCS concentrations (bias = 260 ± 315 cells mL^−1^) than for NES concentrations (bias = 1276 ± 1089 cells mL^−1^). Agreement of mixotroph percentage was also tighter in the CCS (bias = 9 ± 8%) than in the NES (bias = 60 ± 11%). These patterns indicate LysoTracker overestimation, FLP underestimation, or both, particularly in the NES.

### CCS incubation experiment

During a simulated bloom in incubated CCS water, there was high (>50%) LysoTracker staining of water at days 7 and 11 (Figure 6a). Community composition at day 7 was overwhelmingly dominated by diatom taxa (Figure 6b), with more than 80% of reads mapping to diatoms (Bacillariophyta). This percentage of reads mapping to diatoms was lower (∼25%) by the day 11 timepoint, when the stimulated bloom had declined (Supplemental Figure 12).

**Figure 6:**
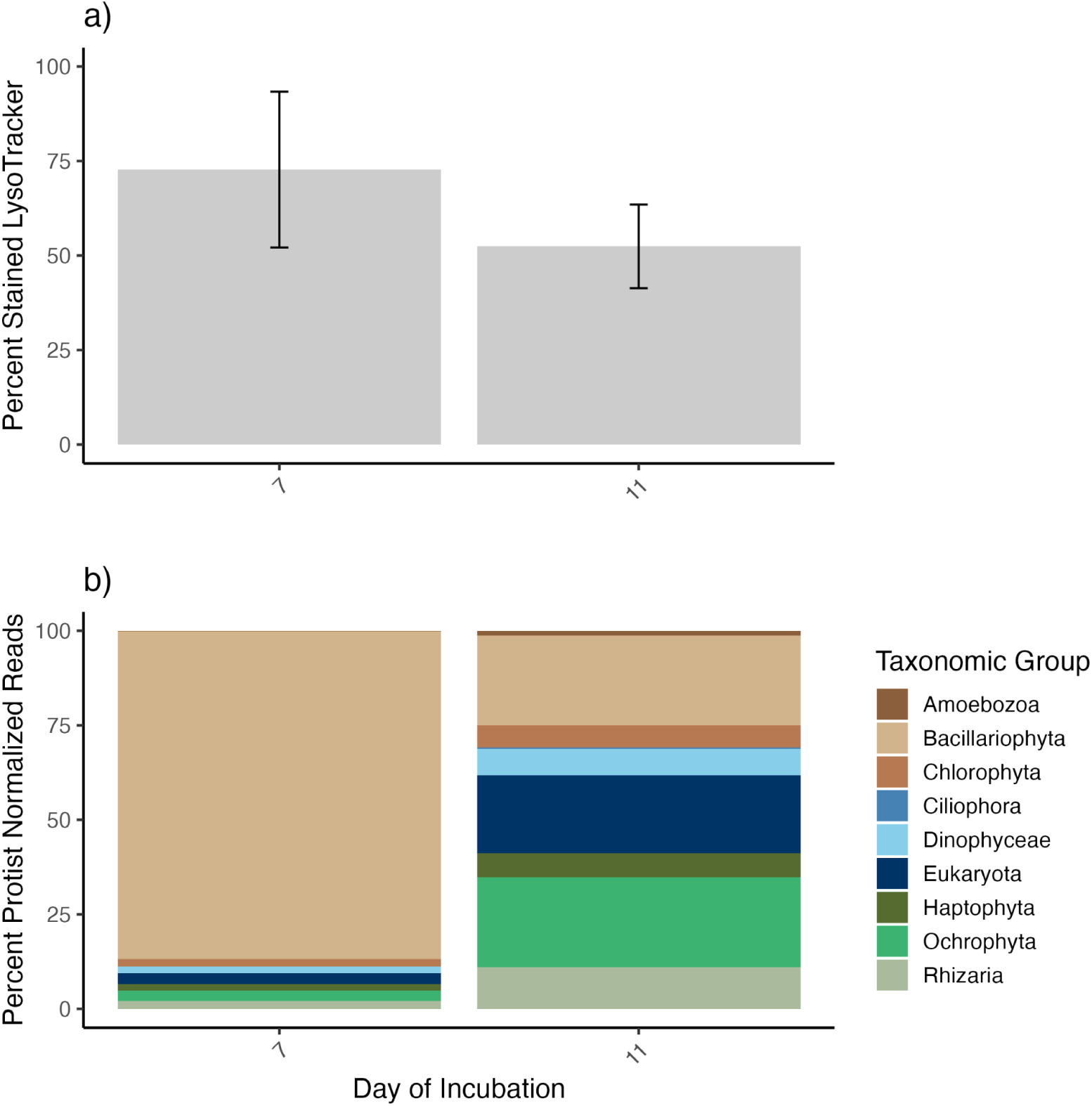
Percent of mixotrophic nanoeukaryotic phototrophs over the 7th and 11th day sampling timepoints in the CCS incubation averaged across biological replicates (a). Taxonomic breakdown of the annotated protist community over these timepoints is displayed (b). Bacillariophyta are diatom taxa while Ochrophyta are non-diatoms.

## Discussion

### Culture-Based insights into acidotropic dye specificity

This study investigated acidotropic dye staining behavior in a diverse assortment of protists in the lab. All diatoms tested demonstrated a high degree of positive staining with LysoTracker (Figure 2; Figure 3) regardless of growth phase and light conditions tested (Supplemental Figure 1), consistent with previous studies that have used acidotropic dyes to measure silica deposition and frustule formation in diatoms (Desclés et al., 2008; McNair et al., 2018; Shimizu et al., 2001). We also found high degrees of positive staining in the diatom *Phaeodactylum tricornutum* cultured in silica-free media, where active silicification is expected to be minimal (Lewin et al., 1958; Martino et al., 2007). This suggests that the observed staining may instead reflect acidic organelles involved in core cellular processes associated with lysosome-like compartments such as autophagy, protein degradation, photosynthetic processes, or other metabolic processes (Grabe & Oster, 2001; Koester et al., 2025; Wilken et al., 2019). The fusiform morphotype observed in *P. tricornutum* throughout this study, typically associated with nutrient rich growth, may indicate elevated metabolic activity and potential for acidic organelle development (Martino et al., 2007). Similarly, other fast-growing diatoms, such as *Chaetoceros spp.* and *Odontella spp.,* stained intensely such that they would be classified as positively stained regardless of the gating thresholds applied (Figure 3; Supplemental Figure 2).

Similarly, the two coccolithophores present in the study, *Gephyrocapsa oceanica* and *Gephyrocapsa huxleyi*, demonstrated more than 50% of cells positively stained with LysoTracker (Figure 2a). Coccolithophores have been demonstrated to possess mixotrophic abilities, and certain species maintain acidic vacuoles during exponential growth that are used for both autophagy (Koester et al., 2025) and phagotrophy (Godrijan et al., 2020; Ye et al., 2024). However, coccolithophores may also have acidic golgi-derived vesicles where coccolith formation occurs as acidic conditions stabilize calcium and carbonate (Brownlee et al., 2015; Marsh, 2003). Therefore, while phagotrophy may have been the cause for staining in the coccolithophores, biomineralization, autophagy, or a combination of these processes may also have contributed.

LysoTracker and LysoSensor demonstrated differences in staining behavior among the isolates. Of the ten diatoms staining positively with LysoTracker, only six stained with LysoSensor, and generally to a lesser extent (Figure 2; Supplemental Table 1). This difference likely reflects the distinct properties of the dyes. LysoSensor, which has a defined acid dissociation constant (pKa 5.1), requires strongly acidic compartments to measurably fluoresce, whereas LysoTracker can label acidic vacuoles of unknown and likely broader pH ranges (Molecular Probes, Invitrogen). As a result, LysoSensor may be more selective, emitting weak or no signal in compartments that are not sufficiently acidic, but more efficient at detecting mature or strongly acidified vacuoles (Huotari & Helenius, 2011). The dyes also use different fluorophores, which may influence their apparent specificity. Green autofluorescence is more common in cells and may contribute to false positives with LysoTracker, while blue autofluorescence is minimal and not likely to interfere with LysoSensor (Tang & Dobbs, 2007).

In contrast to the diatoms, the green algae *Chlamydomonas sp.* (chlorophyte), did not stain with either LysoTracker or LysoSensor and likely represents a true photoautotroph (Figure 2). *Chlamydomonas sp.* has previously been used as a photoautotrophic control in a LysoTracker study and has a low predicted phagotrophy score based on genomic content (Bock et al., 2021; Costa et al., 2022). This culture did not stain with either acidotropic dye, so we hypothesize that this organism is a strict photoautotroph that does not exhibit phagotrophic activity.

*Micromonas polaris* (CCMP 2099) was also initially included in the experiment as a photoautotrophic control. Although one study reported phagotrophy in this species using FLP-based approaches (McKie-Krisberg & Sanders, 2014), subsequent investigations showed no ingestion (Jimenez et al., 2021; Wilken et al., 2019). Genomic models have furthermore predicted photoautotrophy in *M. polaris* (Jimenez et al., 2021) and another species in this genus, *Micromonas pusilla* (Bock et al., 2021). However, a recent transcriptomic model suggested the potential for mixotrophic growth in *M. polaris* (Thomas et al., 2025), adding uncertainty to the trophic mode assignment for this organism. In addition, previous work has shown that the species *M. pusilla* stains with LysoTracker (Rose et al., 2004), while *M. polaris* does not stain with LysoSensor (Jimenez et al., 2021). The latter is consistent with our observation, but this may be due to LysoSensor’s tendency to underrepresent mixotrophy, as demonstrated with confirmed mixotrophs (Figure 2). It is possible our positive LysoTracker staining reflects true mixotrophic activity, but we cannot rule out non-digestive acidic cellular structures in *M. polaris.* Future ingestion experiments using a combination of prey types and sizes are needed to clarify phagotrophic capacity in the *Micromonas* genus.

More than 50% of the populations of two chlorophytes, *Tetraselmis sp.* and *Tetraselmis chui*, stained positively with LysoTracker (Figure 2a). These organisms were initially also included in the study as photoautotrophic controls, as they were previously predicted to have low potential for phagotrophy based on genomic content (Bock et al., 2021). However, grazing ability was assessed through both prey removal experiments (Frost, 1972; Millette et al., 2017) and FLP incubations, and both species were observed to consume prey, albeit in small amounts (Supplemental Figure 4). Since multiple lines of evidence point to *Tetraselmis spp.* being mixotrophic, genomic models trained on a subset of reference species may fail to capture forms of mixotrophic metabolism. This reinforces the need to assess mixotrophic potential through multiple methods.

Three previously characterized mixotrophs were also evaluated. These mixotrophs have been demonstrated to consume prey in light and nutrient replete conditions (McKie-Krisberg et al., 2015). All three demonstrated high staining with LysoTracker, contrasting the LysoSensor results where only *Geminigera cryophila* stained (Figure 2). All three of these mixotrophs consume prey under these conditions (Supplemental Figure 3), which indicates that LysoSensor may fail to identify certain mixotrophs. Under-detection of mixotrophs may be related to cell size, as *G. cryophila* exhibits a larger overall cell volume and presumably larger vacuoles, which may enhance dye accumulation and fluorescence signal.

Within the five confirmed mixotrophic cultures tested in this study, the percent of positively stained cells did not track with ingestion rates. *G. cryophila* demonstrated the highest positive staining of all five mixotrophs with both LysoTracker and LysoSensor (Figure 2). However, the ingestion rate of *G. cryophila* was comparable and within error to the other four species (Supplemental Figures 3-4). Similarly, *Pyramimonas tychotreta* demonstrated the lowest grazing rates of all five mixotrophs, yet demonstrated a higher percent staining with LysoTracker than the *Tetraselmis spp.* and *Mantoniella antarctica.* This agrees with previous observations that acidotropic dyes can be used qualitatively, but do not scale with ingestion rates (Costa et al., 2022). In addition, coccolithophores have been shown to maintain acidic vacuoles regardless of feeding activity, demonstrating the potential for decoupling of acidic vacuoles and actual ingestion (Koester et al., 2025). As acidotropic dyes were unable to track with ingestion rates, we maintain they should instead be used as a qualitative check of whether mixotrophs are potentially important in an ecosystem. For example, a low percentage of LysoTracker-positive nanoeukaryotes likely indicates low levels of mixotrophs and mixotrophy in a system.

We found that cell size plays a role in dye staining interpretations, and warrants size-specific gating thresholds. The large diatoms *Chaetoceros sp. 02* and *Odontella spp.* exhibited high baseline green fluorescence that exceeded the staining gate threshold even before dye addition, requiring a modified gate for these organisms (Figure 3; Supplemental Figure 2). Conversely, the picoeukaryote *M. polaris* demonstrated large increases in green fluorescence relative to its small size (Figure 3), but overall green fluorescence was low (Supplemental Table 4) and it would have been overlooked as unstained using the nanoeukaryote gates (Supplemental Figure 2). Separate gates for picoeukaryotes (Selph et al., 2025) and large cells, or the exclusion of these groups from gating of stained cells as performed in our field study (∼3-20 μm range), can help ensure interpretations are comparable across samples. While taxon-specific thresholds could be established for all cultures in this study, it is necessary to maintain consistency in gating thresholds as gates in the field cannot easily be adjusted in real time due to populations consisting of unknown diversity.

These acidotropic dye culture experiments highlight the complexity of using acidotropic dye staining as a proxy for mixotroph presence. While the goal of applying acidotropic dyes is to detect acidic organelles associated with phagocytosis, these lab results show that biomineralization processes, photosynthetic processes, and autophagy may produce acidic compartments that stain as strongly, or even more strongly, than phagocytic structures. Cell size also should be taken into account when using acidotropic dyes, with gates applied to similarly sized cells. LysoSensor Blue, while requiring a specialized violet laser equipped flow cytometer, may offer a more conservative alternative to LysoTracker, yet still produced false positives in culture and failed to stain known mixotrophs. Additionally, two organisms of the *Tetraselmis* genus, previously considered photoautotrophic, exhibited staining with acidotropic dyes and active prey ingestion, thus enabling identification of new mixotrophic taxa. In contrast, the absence of staining in *Chlamydomonas sp.* supports its designation as a strict photoautotroph and suggests that non-staining may serve as a marker of photoautotrophy in culture.

### Field-based insights into acidotropic dye specificity

To contextualize the acidotropic dye culture experiments, two research cruises were conducted in contrasting coastal ecosystems. For simplicity, we refer to these systems in the framework of r- and k-selection, which describes ecological strategies based on reproductive and resource-use dynamics (Odum, 1969; Parry, 1981). The California Current System (CCS), an active upwelling region, represents an r-selected system where frequent upwelling-driven nutrient inputs selects for fast-growing organisms such as diatoms (Abdala et al., 2022; Closset et al., 2021; Dell’Aquila et al., 2017; Liu et al., 2024). In contrast, the North East Shelf (NES) typifies a k-selected system, characterized by less frequent and more variable nutrient inputs that support more complex trophic structures (Marrec et al., 2021; Yang et al., 2025). In both systems, we performed FLP incubations and staining with the acidotropic LysoTracker to compare these two approaches for detecting mixotrophs *in situ*.

As expected, the summertime NES shelf system exhibited higher concentrations of heterotrophic and mixotrophic nanoeukaryotes as well as bacterial populations representing potential prey compared to the CCS, consistent with a k-selected system. Phototrophic nanoeukaryote and *Synechococcus* abundance was highest nearshore in the NES, reflecting nutrient enrichment from estuarine inputs (Marrec et al., 2021) (Figure 4 a, d). Offshore, phototrophs and *Synechococcus* were most abundant in the subsurface chlorophyll maximum (SCM), while heterotrophic nanoeukaryotes and bacteria remained relatively abundant along the entire transect at all depths (Figure 4). In contrast, the CCS demonstrated lower overall abundance of *Synechococcus* and heterotrophs, with phototrophic abundance more variable across the transect, influenced by the sampling of freshly upwelled versus aged water. The CCS water column was also more vertically mixed, resulting in relatively even abundance distribution across sampled depths. A clear contrast between the systems was observed, as the NES supported higher concentrations of prey and heterotrophic organisms relative to the CCS. Additionally, the nanoeukaryotic community was qualitatively different based on flow cytometry images, with the NES displaying a diversity of size and shapes in the community and noticeable dinoflagellates, while the CCS was primarily dominated by chain forming diatoms (Figure 5 e, f). Based on these dynamics and previous modeling predictions, we would expect a high prevalence of mixotrophs in the k-selective NES relative to the r-selective CCS (Mitra et al., 2014).

Based on the acidotropic dye culture experiments, false mixotroph positives would be expected in the CCS, as much of the community composition was dominated by *Chaetoceros spp.* and other chain-forming diatoms (Supplemental Figure 13; Figure 5 e, f) and all *Chaetoceros* strains isolated from the CCS stained with LysoTracker in culture (Figure 2). However, field data did not support this (Figure 5). LysoTracker staining revealed a greater percentage of mixotrophs in the NES (48.9 ± 22.2%) than in the CCS (9.3 ± 6.8%). These patterns were supported by FLP incubation data, which similarly demonstrated a lower percentage of mixotrophs in the CCS surface waters compared to the NES (Figure 5). However, while FLP incubations and LysoTracker both indicated more mixotrophs in the NES compared to the CCS, these two methods do not demonstrate similar trends within cruise tracks.

FLP incubation results demonstrated that the percent of mixotrophs increased offshore compared to inshore in the NES, while LysoTracker results demonstrated lower percentages of mixotrophy in the offshore stations (Figure 5). Within the NES during the summer months, an observed decoupling of phytoplankton growth and zooplankton grazing has been observed and potentially attributed to mixotrophy in nutrient poor waters (Marrec et al., 2021). Therefore, we hypothesize that mixotrophy may be the most beneficial within the nutrient poor offshore waters of the NES (Ewton, 2025; Marrec et al., 2021; Mitra et al., 2014; Stoecker et al., 2017). FLP incubation data is more consistent with this hypothesis, and therefore may more accurately represent mixotrophic grazing activity within the NES.

Within the CCS, mixotrophic grazing would be expected to track with nitrate or iron dynamics, as these are key limiting nutrients in these waters and play a critical role in photosynthesis (Moore et al., 2013; Till et al., 2019). In this system, water upwelling along narrow shelves has little time to interact with iron rich sediments, resulting in iron limited waters. However, varying bathymetry and deep-water composition create a ‘mosaic of iron limitation’, influencing phytoplankton distributions within the CCS (Till et al., 2019). Along the transect, there was evidence of this iron mosaic, with indications of altered photophysiology caused by iron limitation in the southern stations (5-10) compared to the northern stations (1-4) (Sezinger et al., 2025) where diatoms were in lower relative abundances (Supplemental Fig. 13). Due to suspected iron limitation in the southern stations, mixotrophy could potentially be higher in the south compared to the north as mixotrophs may gain iron through phagotrophy (Maranger et al., 1998). However, FLP experiments do not support this. FLP incubations may have been insufficient to measure detectable mixotroph grazing, as five or less grazers were found per 300 cells surveyed in all CCS microscopy slides (Figure 5). These discrepancies highlight the complexity of measuring mixotroph grazing and presence *in situ* and attributing measurements to environmental factors, as physiological state, community composition, and methodology can modulate the detectability of mixotrophy.

CCS seawater was incubated onboard the research vessel in order to simulate a bloom as part of a larger mesocosm study (Speciale, 2025). Incubation resulted in substantially increased positive staining with LysoTracker, contrasting the low LysoTracker staining values observed across the CCS transect (Figure 6a). The incubation reached peak biomass at day 7, with nanoeukaryote concentrations at a maximum and a small amount of nitrogen still available for growth, indicating the community was likely still in exponential growth (Supplemental Figure 12). At this incubation peak, the incubated community demonstrated high positive staining with LysoTracker, yet was overwhelmingly dominated by diatoms based on RNA taxonomy of the organisms that could be annotated (Figure 6; Supplemental Figure 8). The high levels of staining observed both in isolated diatom cultures and in the natural seawater incubations suggest that prolonged incubation as well as exponential growth may alter cellular physiology, potentially increasing acidification of cellular compartments and susceptibility to LysoTracker staining.

### Comparison of methods and recommendations

The discrepancies observed in mixotroph presence measured by FLP incubations and LysoTracker are likely due to their inherently different targeted mechanisms. Acidotropic dyes label digestive vacuoles, which in dinoflagellates and ciliates can persist from 20 minutes to 3 hours depending on prey ingestion kinetics (Jeong et al., 2010; Sherr et al., 1988). Digestive vacuoles have also been demonstrated to be maintained beyond periods of active ingestion and used for autophagy in addition to phagotrophy (Koester et al., 2025). This complexity complicates interpretations but does not preclude their utility in the field. Acidotropic dyes are reasonable for fieldwork, requiring a brief 10 minute incubation prior to analysis, enabling near-real time estimates of potential mixotrophs following live flow cytometry runs or epifluorescent microscopy. But, as previously discussed, limitations of LysoTracker are that it does not scale with grazing rates (Costa et al., 2022), and may yield false positives (Figure 2).

In contrast, FLP ingestion is a true measure of phagotrophic activity, offering a biogeochemical perspective of grazing rates within a system. Preserved samples can also be analyzed later by flow cytometry or microscopy, avoiding the time constraints of live analysis with acidotropic dyes. However, these short-term incubations only capture grazing within a narrow time window and assume prey provided is of suitable size class and composition for the grazers present (Domaizon et al., 2003; Moorthi et al., 2009). Additionally, the analysis method of FLP incubations can impact data, as low mixotroph grazing or low fluorescent features may be difficult to detect with microscopy and flow cytometry may be uninterpretable due to fluorescent prey sticking to the outside of cells. As a result, FLP incubations likely underestimate mixotrophy due to prey selectivity. Reflecting these methodological biases, agreement between these two methods within each cruise was poor (Supplemental Figure 11).

Despite weak correlations between methods within cruises, both methods indicated that mixotroph presence (Figure 5 a, b), mixotrophic proportion of the population (Figure 5 c, d), and grazing rates (Supplemental Fig. 10) were higher in the NES than in the CCS. The CCS exhibited low mixotrophy by both methods despite being dominated by diatoms, indicating low prevalence of false positives in the field. Given the simplicity of acidotropic dye use and the overall concordance between these approaches, acidotropic dyes could serve as a practical initial tool to assess broad mixotrophic potential in regional aquatic systems, though they are likely not sufficient alone to confirm mixotroph presence due to their potential for false positives. Accounting for background fluorescence (Equation 1) may help reduce false positives, but is not entirely avoidable as photoautotrophic diatoms were seen in this study to stain intensely with both acidotropic dyes (Figure 3). FLP incubations are likely a better choice for estimating mixotrophy *in situ* due to their comprehensive estimates of grazing and lower possibility for false positives (Wilken et al., 2019). Taken together, these methods offer complementary insights with acidotropic dyes facilitating rapid screening and FLP incubations providing more confident estimates. Applying both methods in tandem may therefore be an effective strategy for advancing our understanding of mixotroph distributions and their ecological roles across environmental gradients.

### Conclusions

Acidotropic dyes offer a promising approach for detecting mixotrophs *in situ* due to their relative ease of use and potential for near real-time application. In this study, we compared laboratory and field applications to evaluate their utility. Our results show that false positives were observed in diatom species, indicating that LysoTracker staining occurs in these organisms due to acidic processes unrelated to phagotrophy, such as biomineralization, autophagy, or cellular storage. In contrast, one non-biomineralizing photoautotrophic control, *Chlamydomonas sp.,* did not stain with either acidotropic dye tested, supporting its classification as a true photoautotroph. LysoSensor produced fewer false positives than LysoTracker, yet also underestimated mixotrophy, as two of five known mixotrophs did not stain with LysoSensor. In field samples, LysoTracker results broadly aligned with FLP-based grazing estimates at a broad regional scale, with higher mixotroph abundance and mixotrophic activity in the NES compared to the CCS, but diverged along finer coastal gradients, underscoring that the two approaches target different cellular mechanisms. FLP assays measure active ingestion, while acidotropic dyes may better reflect mixotrophic potential. Thus, acidotropic dyes may serve as a useful tool for quickly and qualitatively evaluating mixotrophy. In the field, it is recommended to couple acidotropic dyes with at least one other measurement of phagotrophy, and to consider community composition when assessing the importance of mixotrophy in a given aquatic ecosystem.

## Acknowledgements

We acknowledge support from the National Science Foundation (NSF) Office of Polar Programs (ANT-LIA-2240780 to NRC), NSF Division of Ocean Sciences (OCE-2322676, OCE-1655686 to SMD), NSF CAREER Awards (OCE-1751805 to AM), Alfred P. Sloan Research Fellowship to NRC, Simons Foundation (SFI-LS-ECIAMEE-00006690 to NRC), and University of Georgia’s Skidaway Institute of Oceanography. We thank the scientists and crew on the R/V *Sally Ride* and R/V *Endeavor,* and all scientists that provided culture organisms used in this study: Rebecca Gast, Lucy Quirk, and Mak Saito as well as the NCMA and Milford strain collections. Microscopy data and image data for *Tetraselmis spp.* grazing was produced with instrumentation at the Biomedical Microscopy Core at the University of Georgia. We would like to thank Pierre Marrec (URI) for sharing his flow cytometry expertise on the NES cruise and Skidaway Institute summer interns Neha Shah and Felipe Quintana for their assistance in culturing.

## Author contributions

CCZC and NRC designed the study. CCZC cultured organisms, analyzed lab and field data, and wrote the first draft manuscript under the guidance of NRC. SMD and AM were Chief Scientists on the NES and CCS cruises, respectively, and coordinated fieldwork. EME and SMD contributed to NES sampling efforts and data interpretations. AM and EVS led the CCS incubation experiments and provided taxonomic metatranscriptome information. NCM performed prey removal experiments with laboratory cultures. SW and SS contributed to data interpretations. All authors reviewed and edited the manuscript and approved of the final version.

## Data availability

Data and code used in this analysis are publicly available at https://github.com/CohenLabUGA/AcidotropicDyes. Attune CytPix flow cytometry photos representing the community composition from both cruises can be found at https://zenodo.org/records/17108625.

## Artificial Intelligence statement

The authors declare that generative AI was used to assist in the refinement of code used in this paper. The AI usage was confined to tasks such as coding error identification and optimizing code syntax. All scientific interpretations and conclusions are derived from the authors.

## Supplemental Materials

**Supplemental Figure 1:**
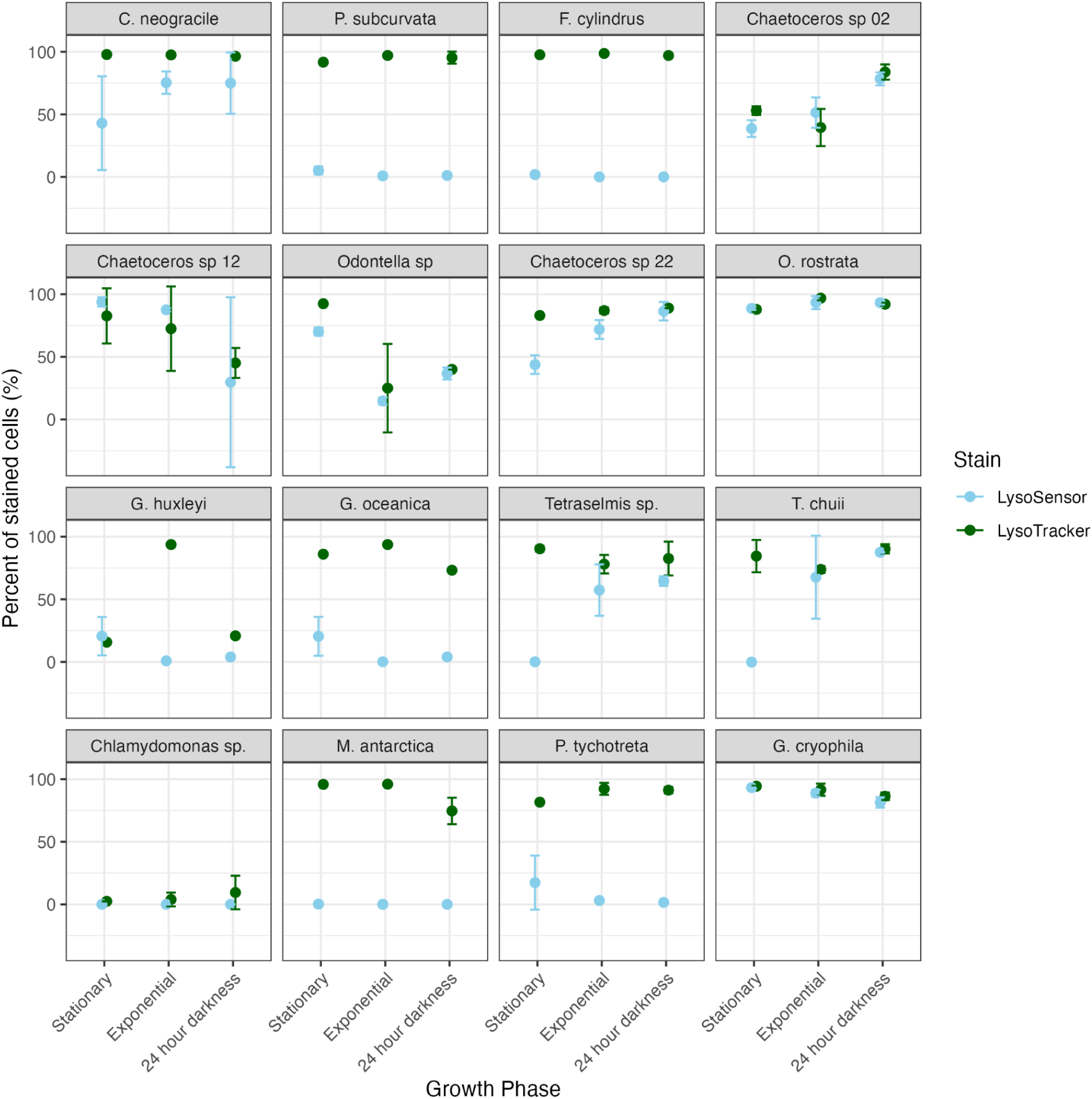
Percent of cells stained by LysoTracker (green) and LysoSensor (blue) across three growth phases (stationary, exponential, and after 24 hours of darkness). Error bars represent the standard deviation of two technical replicates.

**Supplemental Figure 2:**
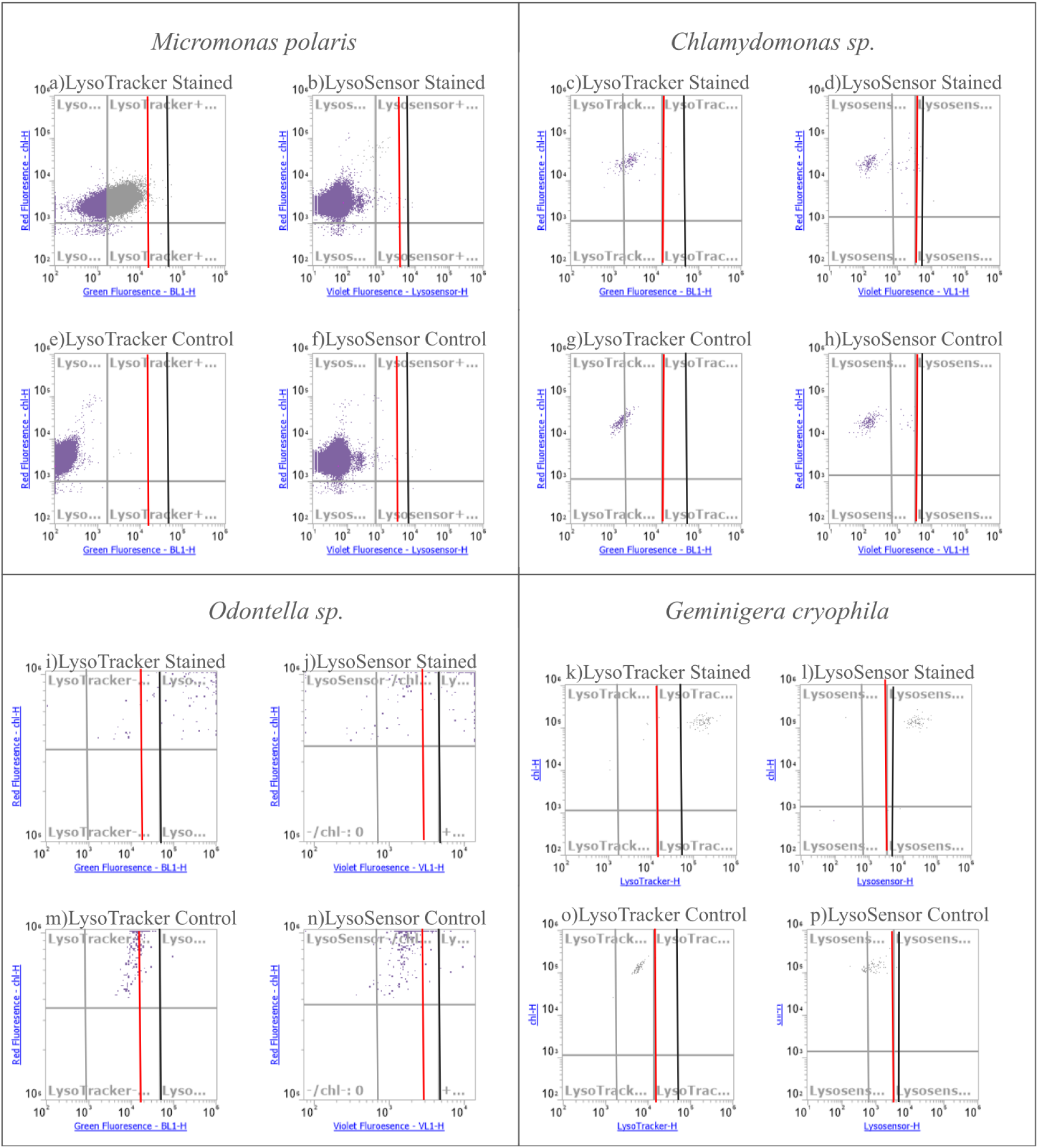
Gates used for determining acidotropic dye staining in cultures. A picoeukaryote specific gate was made for *Micromonas polaris* (a, b, e, f) and is reflected on all plots as the gray line. The standard nanoeukaryote gates are demonstrated by the red line with representative nanoeukaryotes *Chlamydomonas sp.* (c, d, g, h) and *Geminigera cryophila* (k, l, o, p). The gate for large diatoms *Chaetoceros sp. 02* (UNC2302), *Odontella sp.* (UNC2314), and *O. rostrata* (UGA01) is demonstrated with the black line with representative large diatom *Odontella sp.* LysoTracker staining was represented by elevated green fluorescence (a, c, i, k). LysoSensor staining was represented by elevated blue fluorescence (b, d, j, l). The unstained samples (e-h, m-p) were subtracted from their corresponding stained sample.

**Supplemental Table 1:** Number of biological replicates (n) for each organism with each type of stain and flow cytometer combination with reported average and standard deviation of the percent of cells that were positively stained. The reported mean and standard deviation of LysoTracker values in Figure 2 is a combination of biological replicates from both flow cytometers. The range of cells examined refers to the cell concentrations (cells mL^−1^) of the cultures during staining. Note that large cells could not be run on the Guava easyCyte (Table 1).

| Staining Summary ((n); Mean ± StDev) | | | | |
| --- | --- | --- | --- | --- |
| Culture Name | LysoTracker (CytPix) | LysoTracker (Guava) | LysoSensor (CytPix) | Range of Cells Examined |
| C. neogracile | (2); 97.51 ± 1.18 | (2); 98.34 ± 0.00 | (3); 53.57 ± 38.21 | 34390 – 183420 |
| Chaetoceros sp. 02 | (2); 68.05 ± 40.37 | NA | (2); 68.49 ± 24.13 | 80 – 503290 |
| Chaetoceros sp. 12 | (2); 72.50 ± 33.76 | NA | (2); 63.63 ± 33.85 | 1230 – 4010 |
| Chaetoceros sp. 22 | (1); 86.96 ± NA | (3); 85.14 ± 1.54 | (2); 73.58 ± 2.46 | 1820 – 5940 |
| Chlamydomonas sp. | (2); 3.30 ± 0.85 | (3); -0.34 ± 0.29 | (2); 0.60 ± 0.85 | 50 – 13430 |
| F. cylindrus | (1); 98.67 ± NA | (4); 94.17 ± 9.54 | (2); 0.02 ± 0.03 | 2100 – 363120 |
| G. cryophila | (1); 91.60 ± NA | (4); 86.96 ± 7.55 | (2); 92.61 ± 5.34 | 590 – 13950 |
| G. huxleyi | (1); 91.91 ± NA | (3); 63.06 ± 0.40 | (2); 0.66 ± 0.28 | 10050 – 35990 |
| G. oceanica | (1); 93.74 ± NA | (3); 94.03 ± 3.25 | (3); 0.02 ± 0.07 | 9010 – 438220 |
| M. antarctica | (2); 91.46 ± 5.83 | (3); 95.37 ± 2.59 | (3); -0.02 ± 0.10 | 2040 – 23260 |
| M. polaris | (2); 50.87 ± 18.21 | NA | (2); -1.11 ± 1.57 | 130 – 1342160 |
| O. rostrata | (2); 95.90 ± 1.37 | NA | (2); 96.11 ± 3.90 | 860 – 5580 |
| Odontella sp. | (2); 86.18 ± 6.88 | NA | (3); 60.27 ± 42.15 | 1900 – 711960 |
| P. subcurvata | (1); 97.03 ± NA | (4); 95.74 ± 2.31 | (2); 0.48 ± 0.58 | 670 – 6130 |
| P. tricornutum | (3); 83.91 ± 24.46 | NA | (3); 60.98 ± 26.47 | 870 – 371650 |
| P. tychoetreta | (5); 95.10 ± 2.80 | (4); 95.46 ± 1.64 | (5); 15.20 ± 12.74 | 1460 – 16700 |
| T. chui | (1); 88.98 ± NA | (3); 90.68 ± 1.76 | (2); 69.02 ± 1.92 | 460 – 106770 |
| Tetraselmis sp. | (1); 85.19 ± NA | (3); 96.47 ± 4.22 | (2); 49.46 ± 11.21 | 510 – 202460 |

**Supplemental Table 2:** Gain and voltage settings for the channels used in this study for both Guava EasyCyte and Attune CytPix flow cytometers. Detection wavelengths of each channel are also listed.

| Instrument Settings for Guava EasyCyte and Attune CytPix Flow Cytometers |  |  |  |  |  |  |
| --- | --- | --- | --- | --- | --- | --- |
| Channel | Gain and Voltage Settings |  |  |  | Detection Wavelengths |  |
|  | Guava Gain - Culture Tests & NES FLP | Guava Gain - LysoTracker CCS | Guava Gain - LysoTracker NES | CytPix Voltage - Culture Tests | CytPix Wavelength (nm) | Guava Wavelength (nm) |
| FSC | 4.97 | 4 | 4 | 240 | — | — |
| SSC | 2.95 | 1 | 1 | 330 | 480/10 | 488/16 |
| Green | 2.95 | 2.95 | 2.95 | 377 | 530/30 | 512/18 |
| Yellow | 112 | 98.7 | 98.7 | — | — | 575/25 |
| Red | 2.95 | 2.95 | 4 | 280 | 695/40 | 695/50 |
| Violet | — | — | — | 354 | 440/50 | — |
| High acquisition settings | Off | Off | On | — | — | — |

**Supplemental Figure 3:**
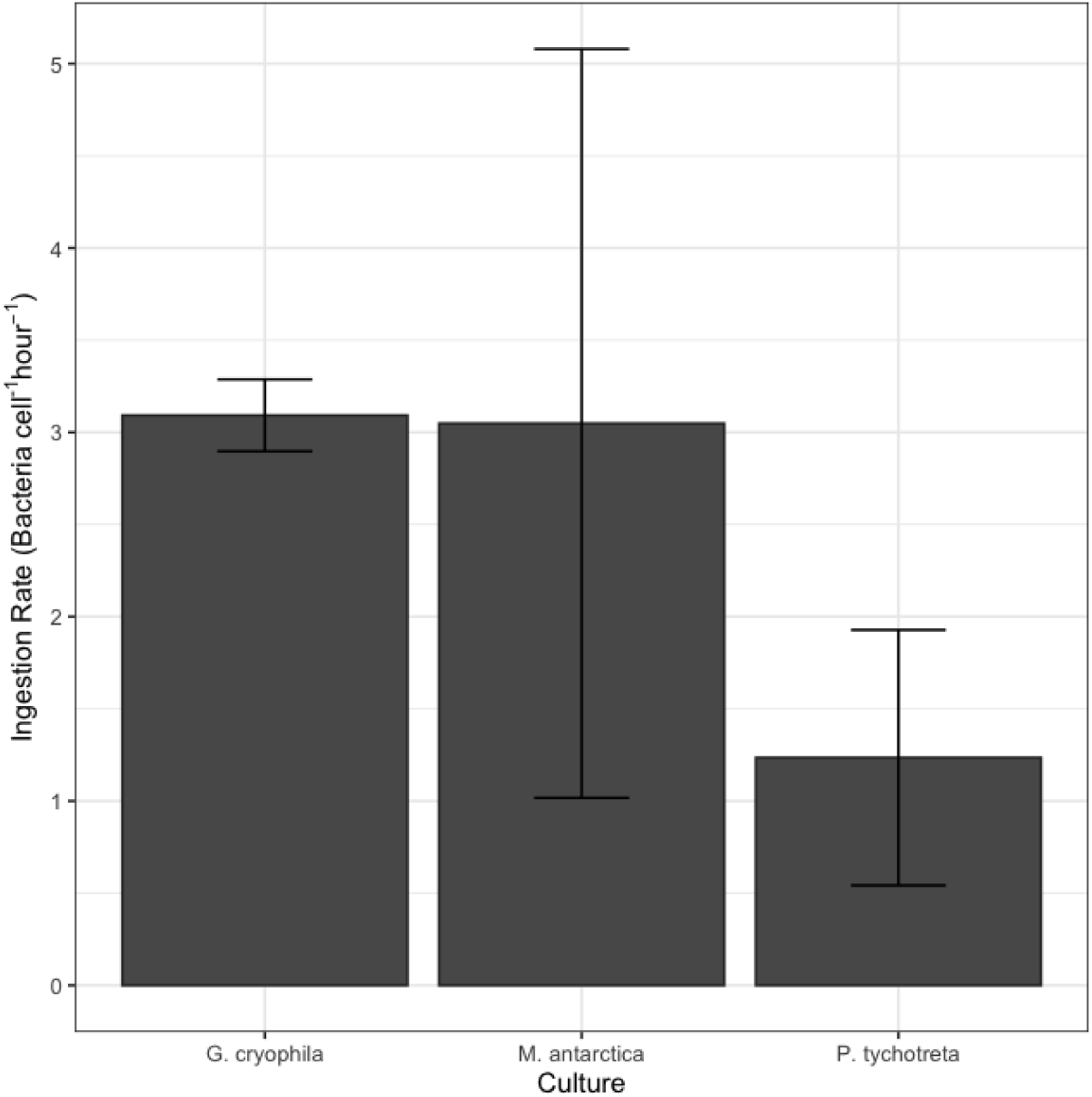
Grazing data for the three Southern Ocean mixotrophs in replete medium and standard light conditions: *G. cryophila, M. antarctica,* and *P. tychotreta.* Ingestion rate (bacteria cell^−1^ hour^−1^) is measured via prey removal experiments. Bars represent the average and error bars represent the standard deviation of biological replicates.

**Supplemental Figure 4:**
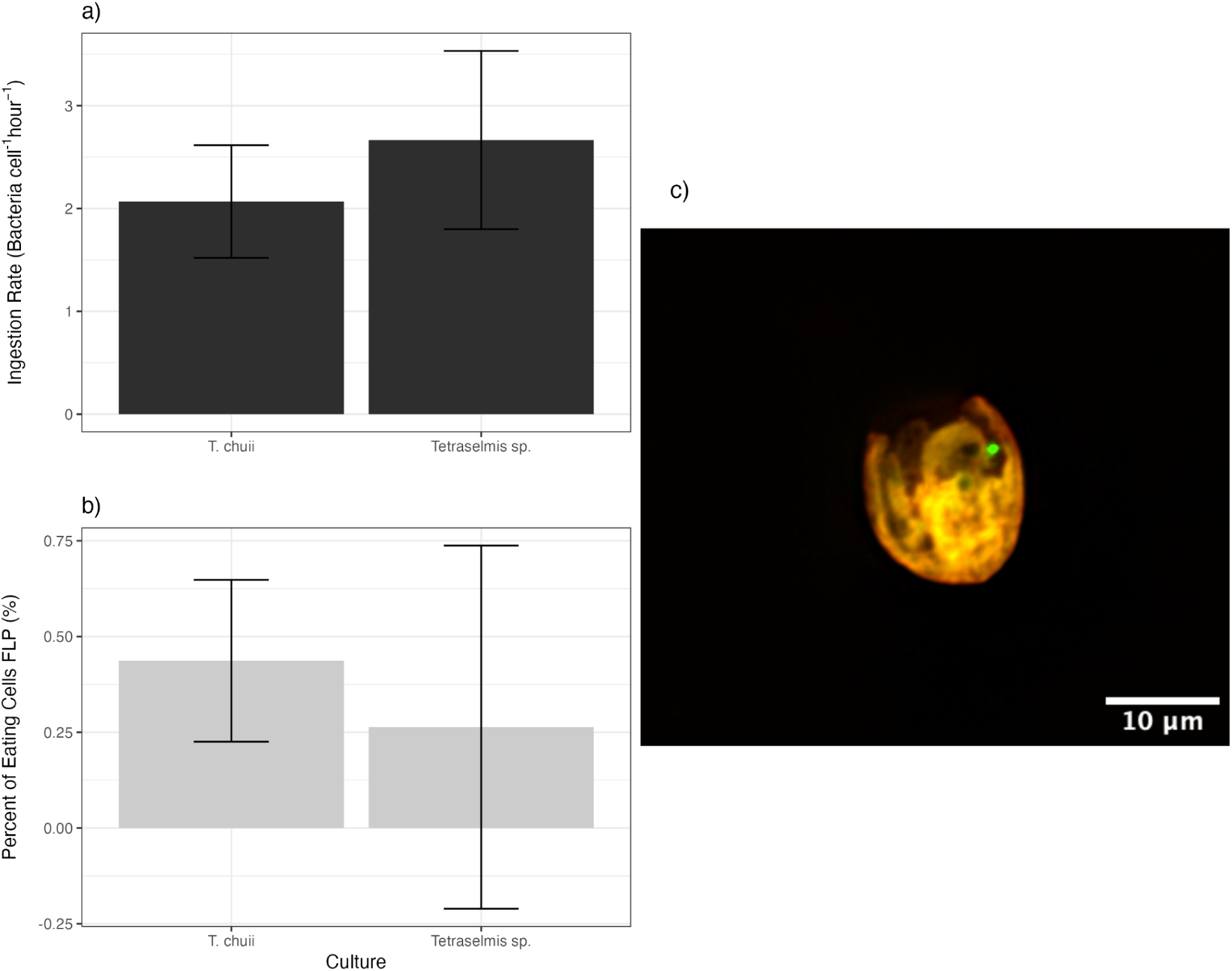
Grazing information for the *Tetraselmis spp.* Grazing was assessed both through prey removal experiments (a) and FLP experiments analyzed using microscopy (b). An example of a mixotrophic *Tetraselmis sp.* cell (c) is displayed. Bars represent the average value across three biological replicates, error bars represent standard deviation.

**Supplemental Figure 5:**
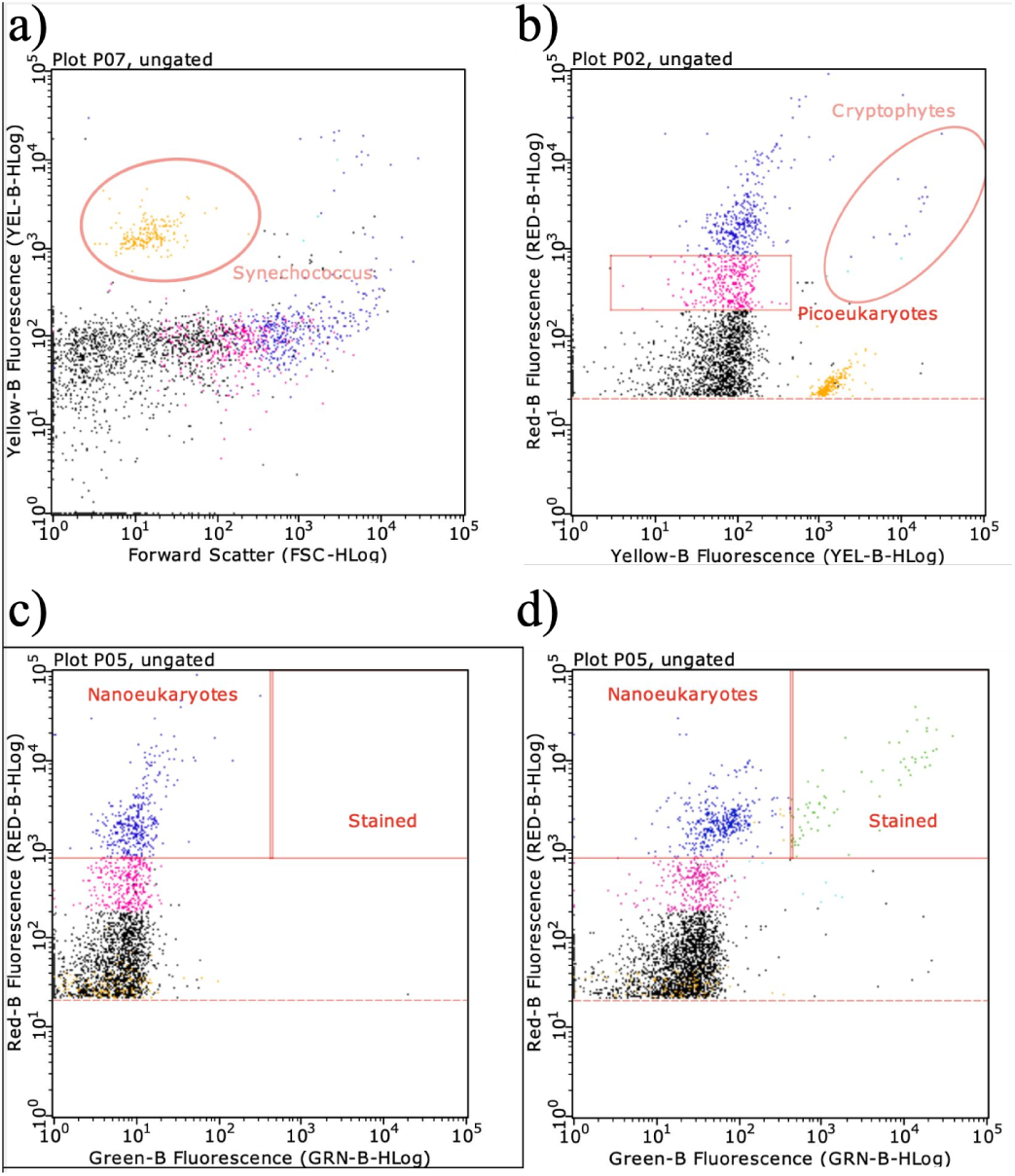
Example gates from LysoTracker staining of field samples using a Guava easyCyte HT flow cytometer. *Synechococcus* is identified based on low FSC and high yellow fluorescence (a). Picoeukaryotes were identified by medium red fluorescence (b). Nanoeukaryotes were identified based on high red fluorescence (c). Mixotrophic cells were identified by high green and red fluorescence (c, d). Stained samples (d) were subtracted from unstained control samples (c).

**Supplemental Figure 6:**
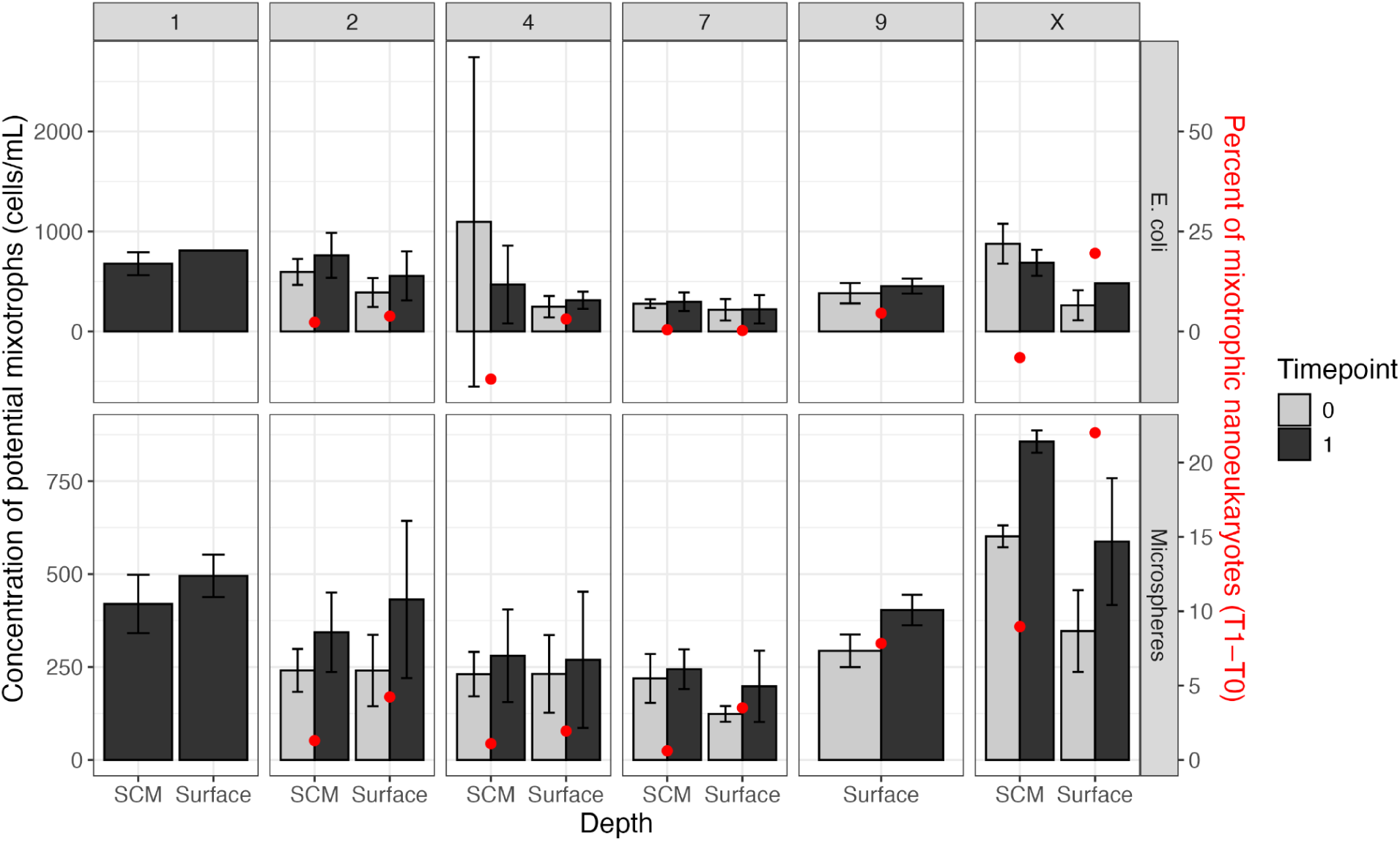
Comparison of prey types (*E. coli* and microspheres) for FLP flow cytometry data in the NES. Left y axis and bars represent the concentration of mixotrophs across the FLP incubation sampling timepoints. Note that timepoint 0 is subtracted from timepoint 1 to determine percent mixotrophs relative to total phototrophic nanoeukaryote in a given sample. Right y axis and red dots represent the overall percent of mixotrophic nanoeukaryotes. Error bars represent the standard deviation of two biological replicates.

**Supplemental Figure 7:**
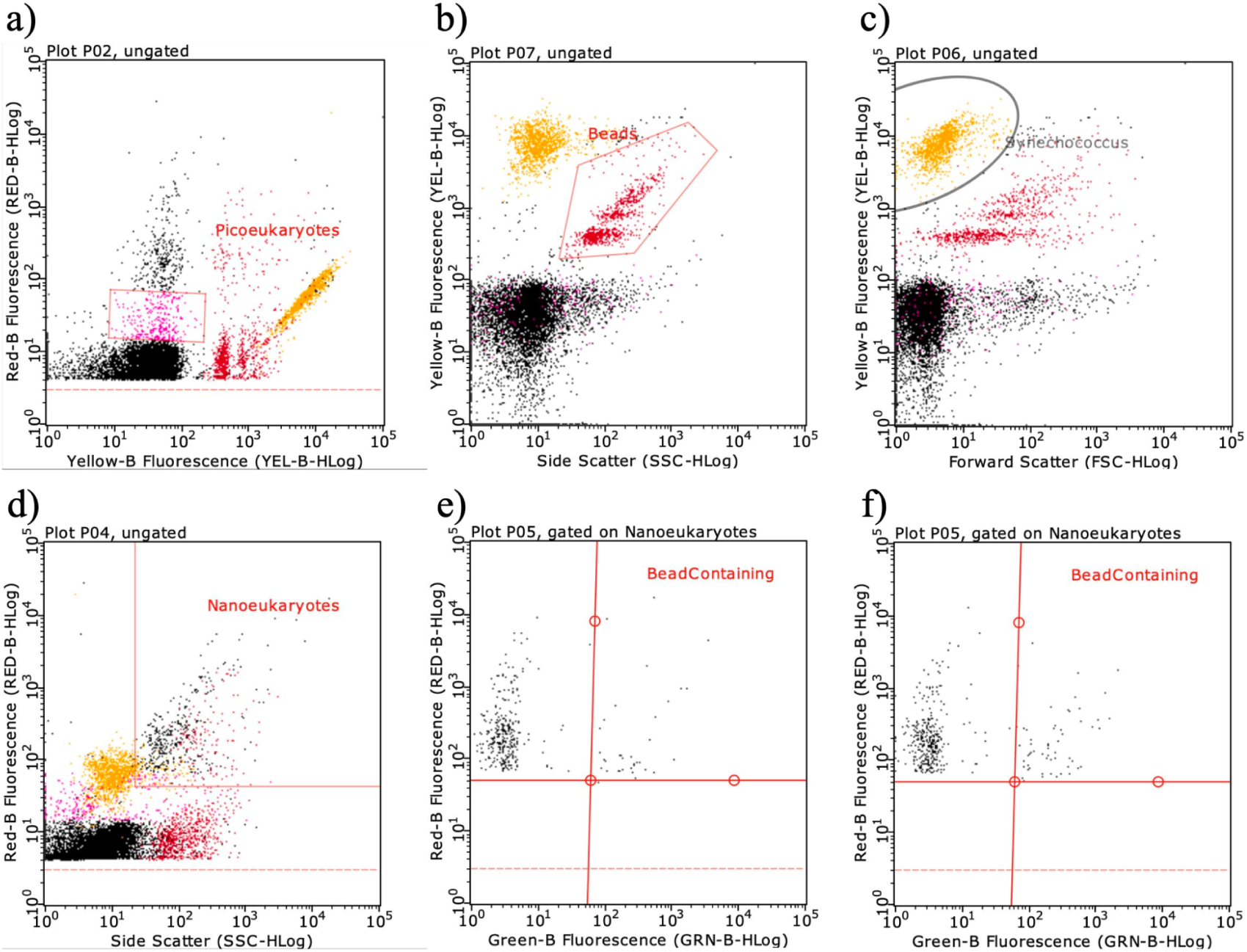
Example gates representing the FLP experiments on the NES cruise. Picoeukaryotes were defined based on medium/low red fluorescence (a). Beads were identified based on low side scatter and high yellow fluorescence to differentiate from *Synechococcus* (b). *Synechococcus* was identified based on high yellow and low FSC (c). Nanoeukaryotes were identified based on high red fluorescence and SSC (d). Finally, mixotrophs were identified based on nanoeukaryotes that had high green fluorescence (e).

**Supplemental Table 3:** Number of biological FLP incubation replicates for microscopy at each station in the NES.

| Station | Biological Replicates<br>Surface | Biological Replicates<br>SCM |
| --- | --- | --- |
| 1 | 1 | 1 |
| 2 | 1 | 1 |
| 4 | 2 | 1 |
| 7 | 1 | 2 |
| 9 | 1 | 2 |
| X | 1 | 1 |

**Supplemental Figure 8:**
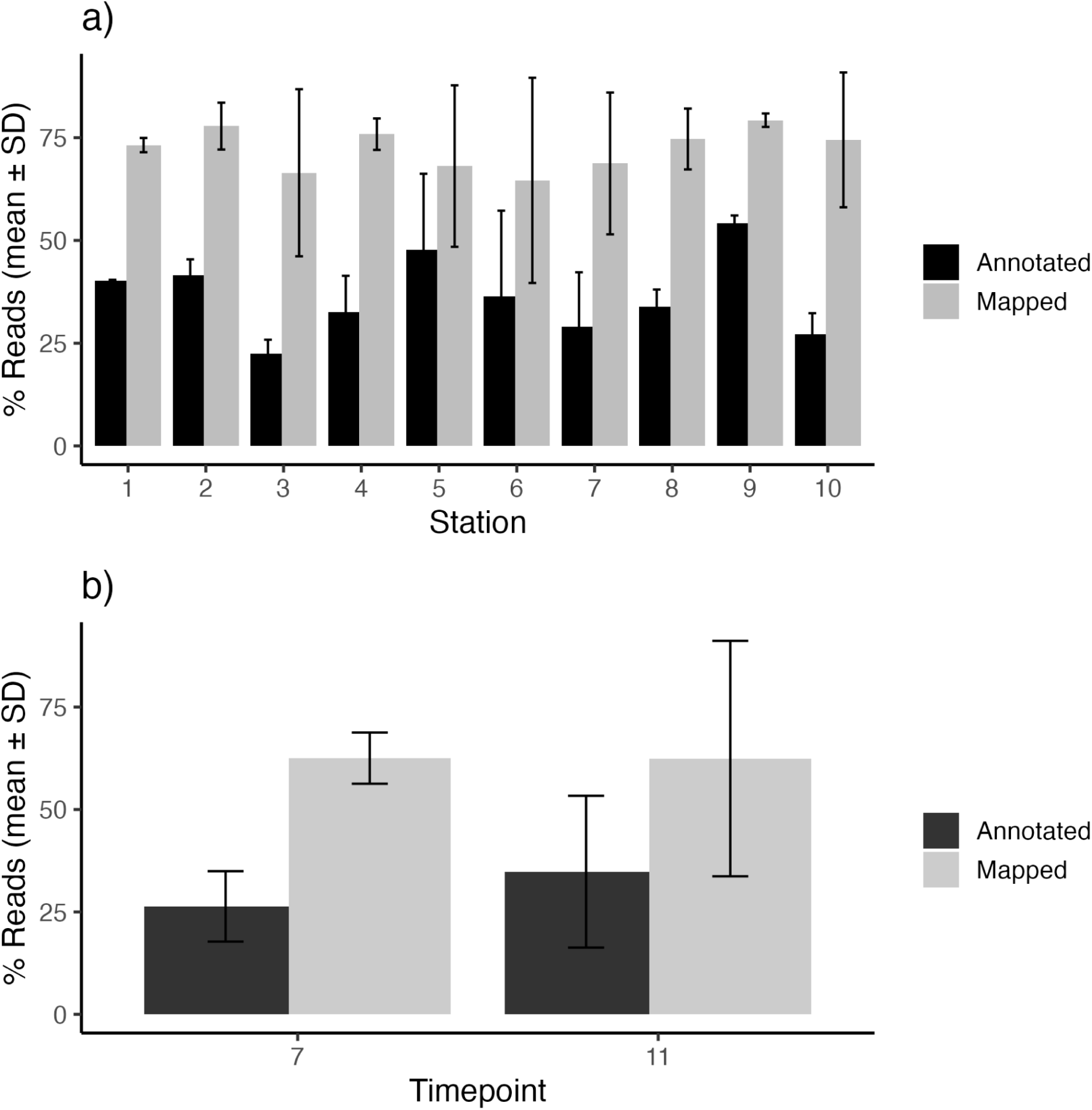
Percent of reads mapped (gray) and annotated (black) from both the CCS stations (a) and incubation (b). Error bars represent the standard deviation of 3 biological replicates.

**Supplemental Figure 9:**
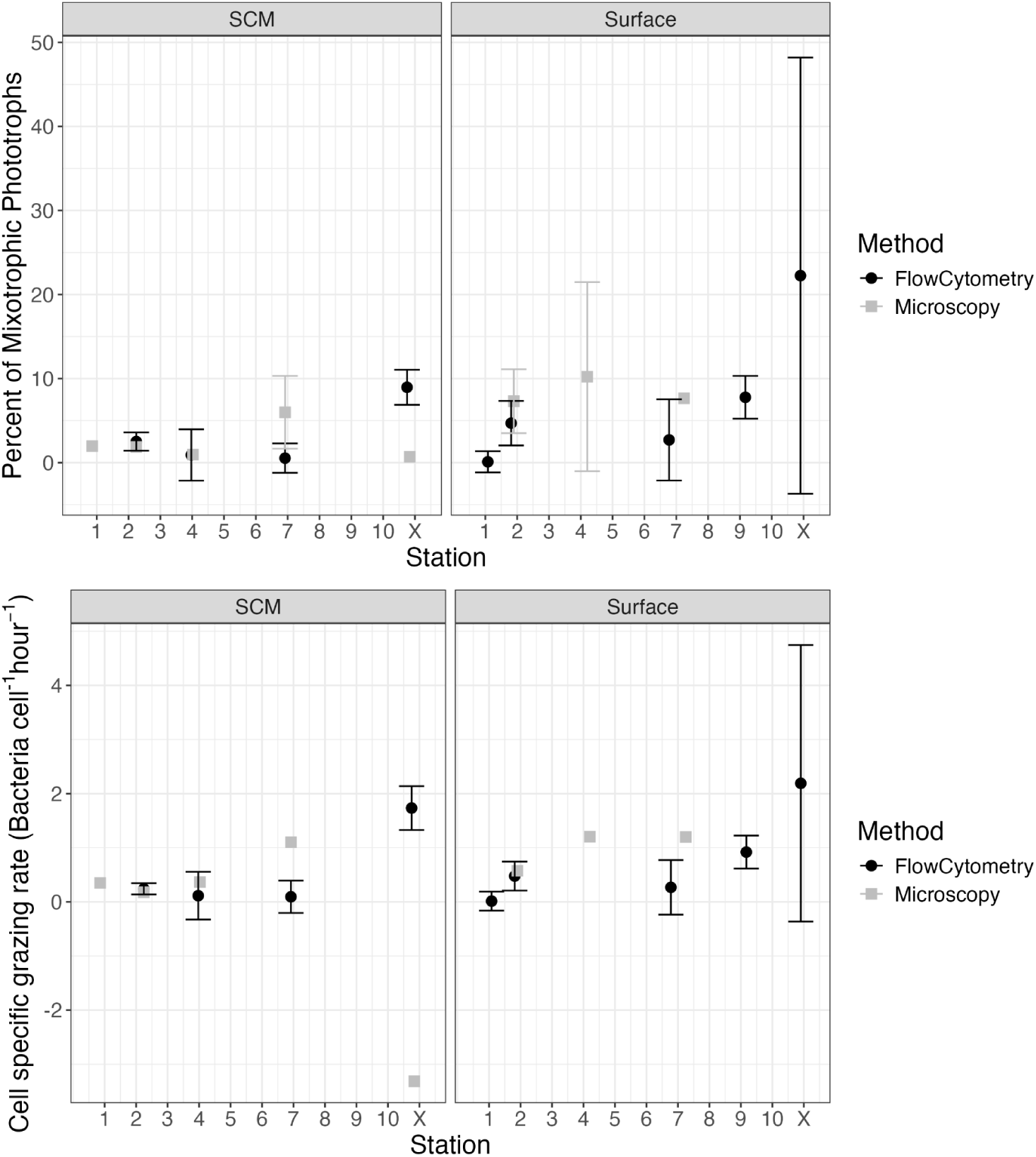
Analysis of percent of mixotrophic protists (top) and cell specific grazing rate (bottom) for the NES. FLP incubation analysis type is represented with color and shape, with microscopy in gray squares and flow cytometry in black circles. Points represent the average across biological replicates with error bars indicating standard deviation.

**Supplemental Figure 10:**
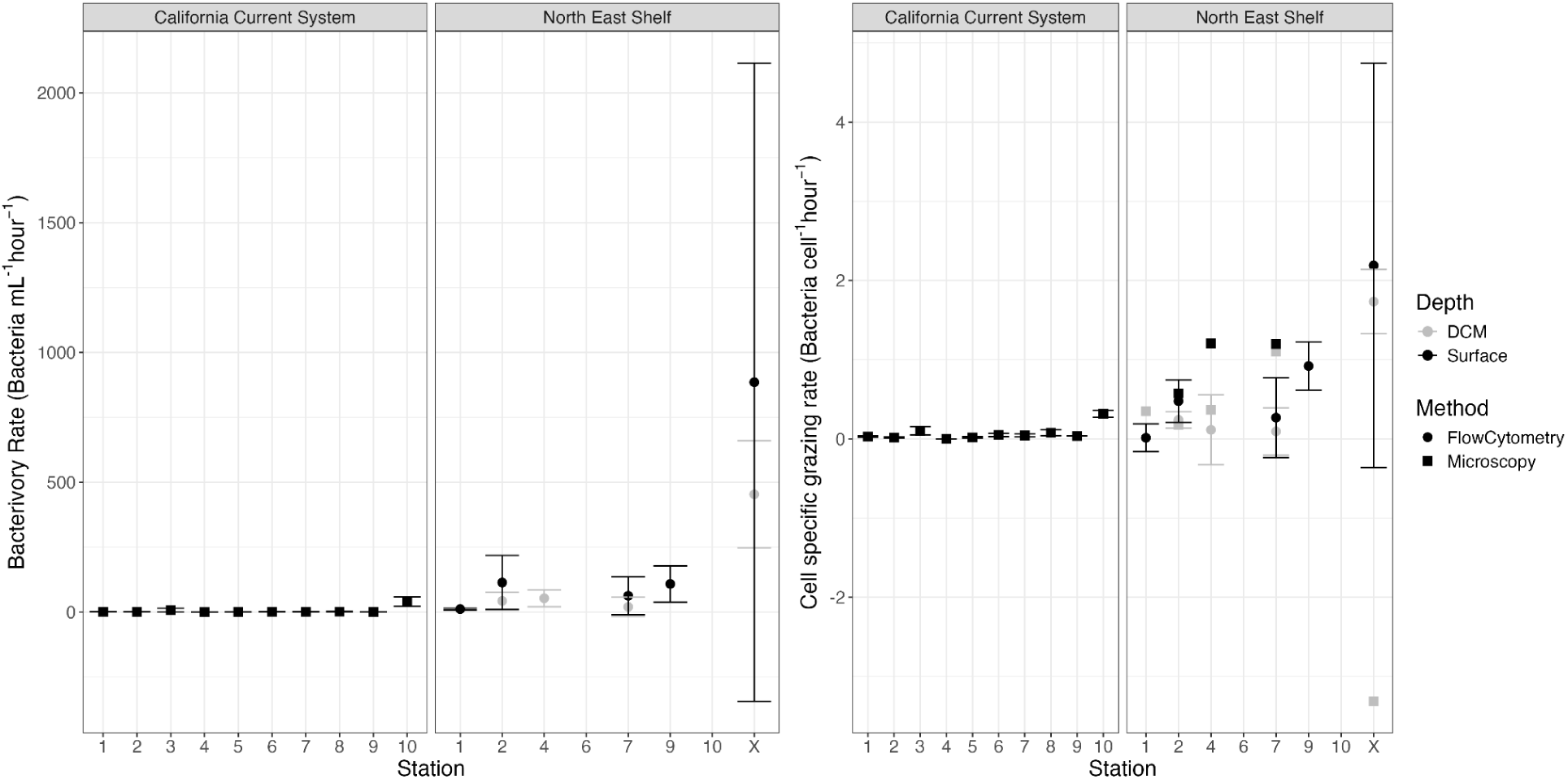
Comparison of community bacterivory rate (left) and grazing rate (right) from the NES and CCS cruises. Dots represent the mean, error bars represent standard deviation of biological replicates. Cell specific grazing rate contains data from both flow cytometry (circles) and microscopy (squares) FLP incubation analysis techniques and depths DCM (gray) and surface (black). All CCS FLP data was collected via microscopy.

**Supplemental Table 4:**
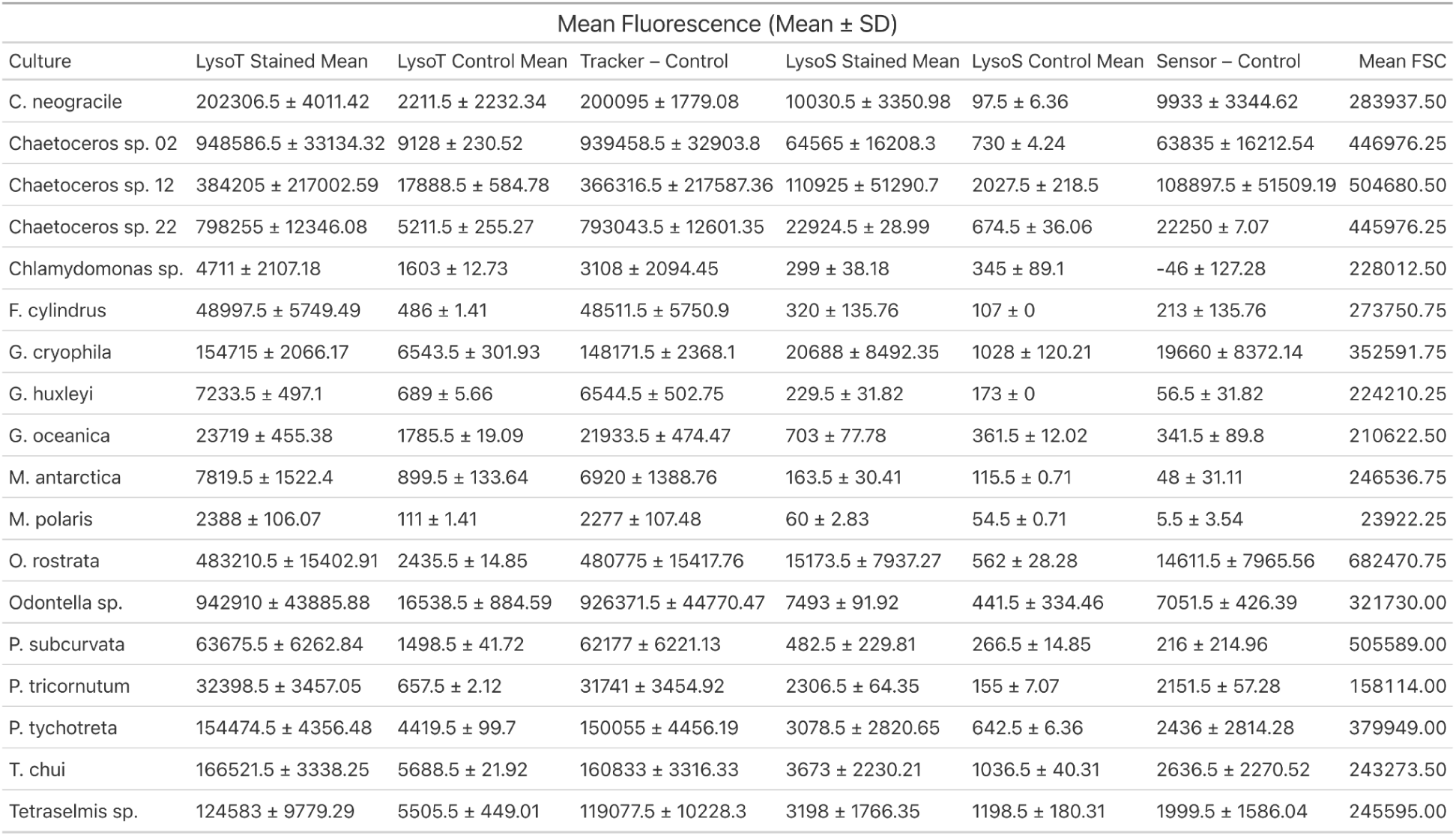
Analysis of different fluorescence parameters between the stained and control samples. Reported is the mean green fluorescence intensity for both LysoTracker (LysoT) stained and control, the difference in the mean fluorescence between the LysoTracker stained and control samples (Tracker - Control), the mean blue fluorescence for both LysoSensor (LysoS) stained and control, the difference in mean blue fluorescence between LysoSensor stained and control samples (Sensor - Control), and the mean forward scatter (FSC) for each culture. Averages and standard deviations are calculated from two technical replicates.

**Supplemental Figure 11:**
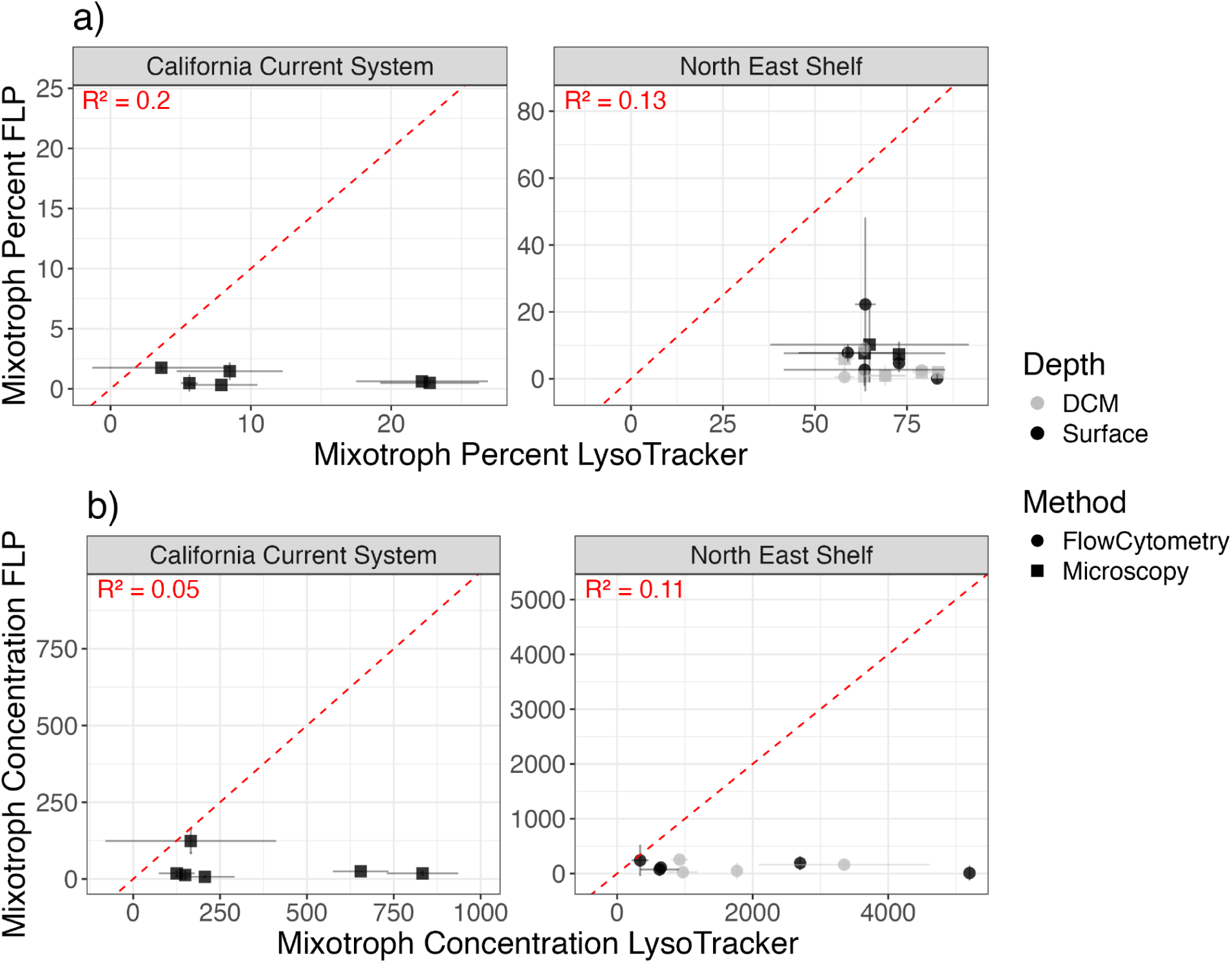
Comparisons between FLP and LysoTracker methods for determining mixotroph percentage (a) and mixotroph concentration (b). Red dotted lines represent a 1:1 line with *R^2^* values displaying relatedness of the measurements. Depth of sample is indicated by color (black surface; gray DCM/SCM) and method of FLP assessment is indicated by shape (square microscopy; circle flow cytometry). Error bars indicate the standard deviation of biological replicates.

**Supplemental Figure 12:**
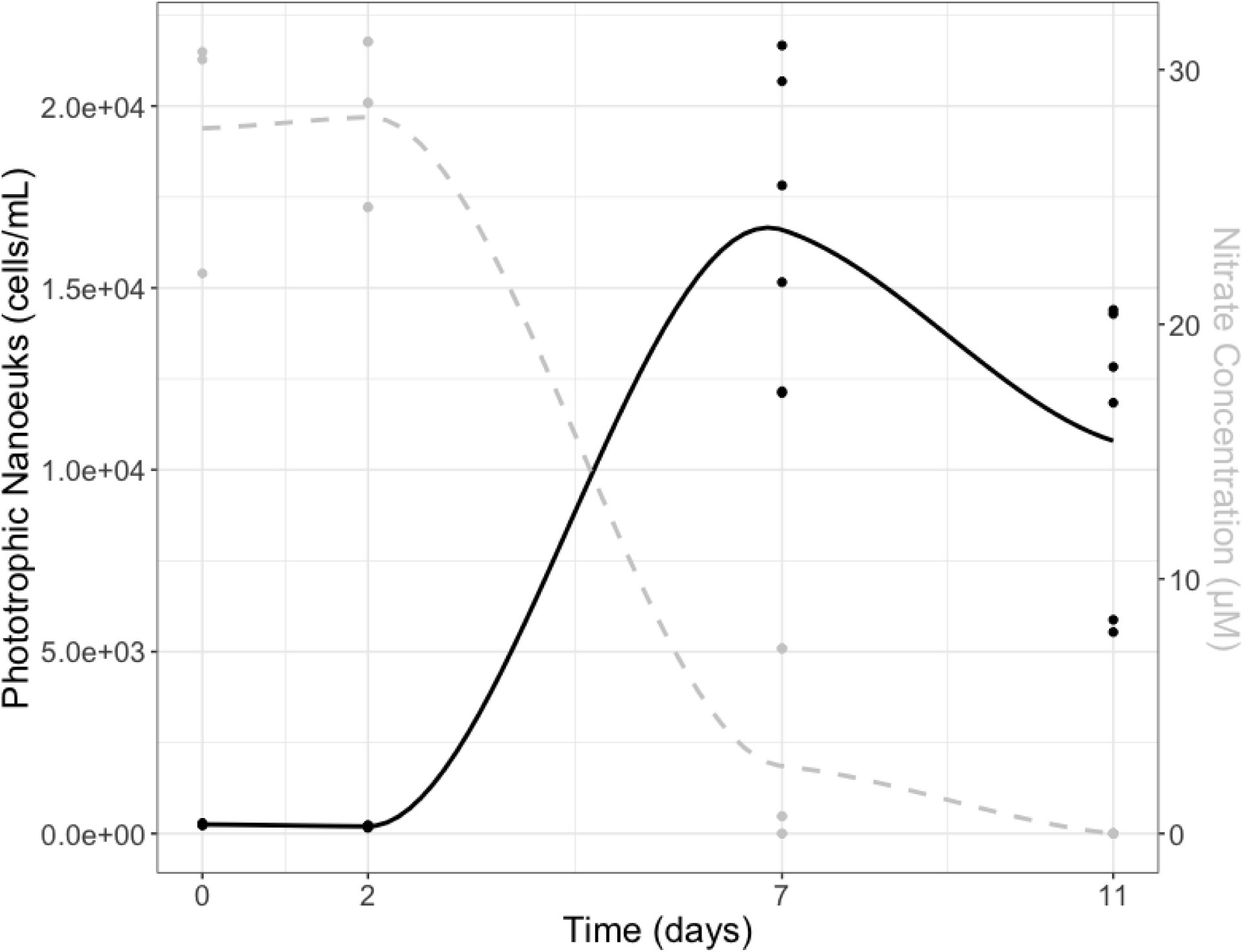
Phototrophic nanoeukaryotes (cells mL^−1^, black) and nitrate concentration (µM, dashed gray) over the course of the CCS incubation experiment.

**Supplemental Figure 13:**
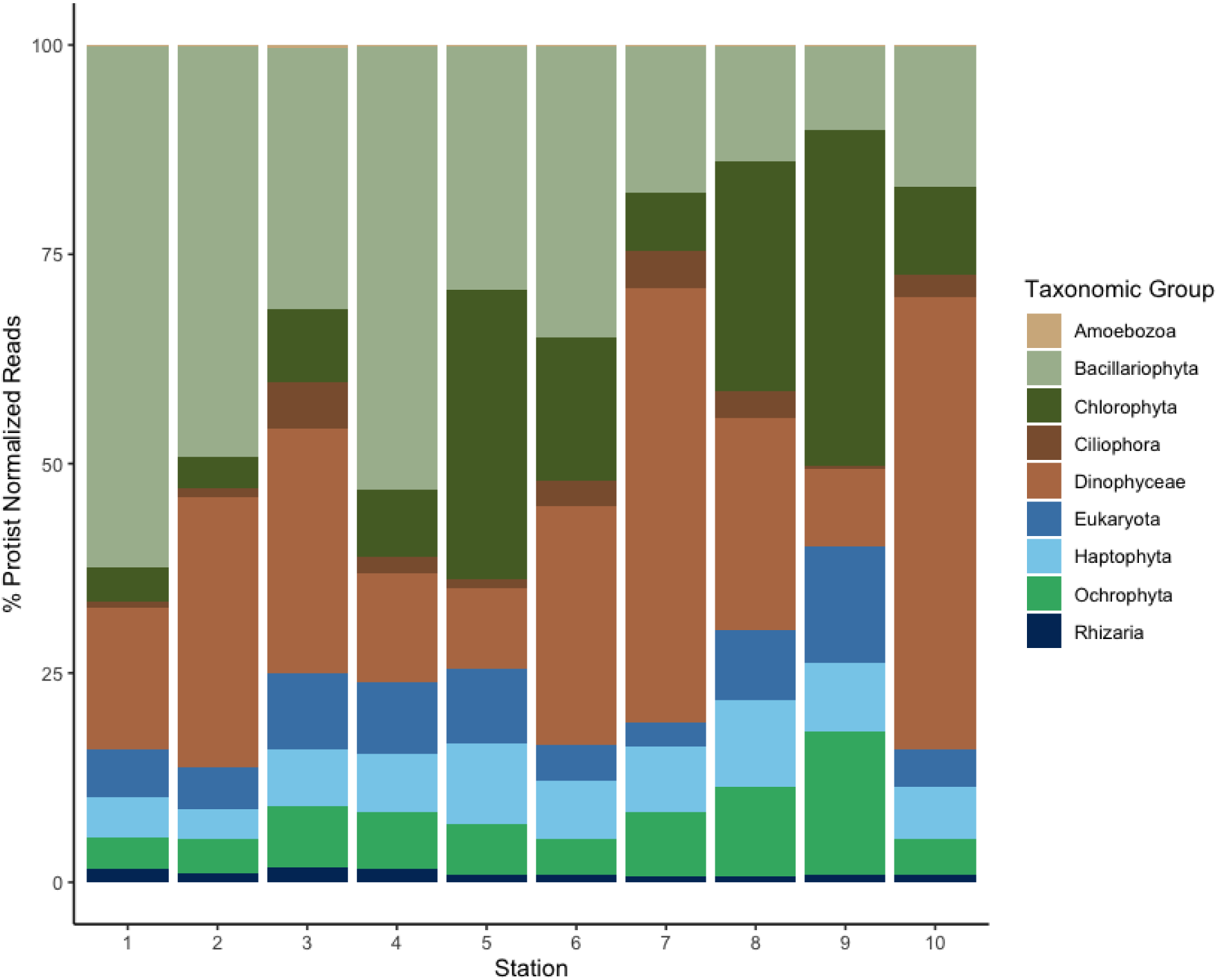
Taxonomic breakdown of the protists present along the CCS transect showing only the annotated fraction.

